# The neuropeptidergic connectome of *C. elegans*

**DOI:** 10.1101/2022.10.30.514396

**Authors:** Lidia Ripoll-Sánchez, Jan Watteyne, HaoSheng Sun, Robert Fernandez, Seth R Taylor, Alexis Weinreb, Mark Hammarlund, David M Miller, Oliver Hobert, Isabel Beets, Petra E Vértes, William R Schafer

## Abstract

Efforts are currently ongoing to map synaptic wiring diagrams or connectomes in order to understand the neural basis of brain function. However, chemical synapses represent only one type of functionally important neuronal connection; in particular, extrasynaptic, “wireless” signaling by neuropeptides is widespread and plays essential roles in all nervous systems. By integrating single-cell anatomical and gene expression datasets with a biochemical analysis of receptor-ligand interactions, we have generated a draft connectome of neuropeptide signaling in the *C. elegans* nervous system. This connectome is characterized by a high connection density, extended signaling cascades, autocrine foci, and a decentralized topology, with a large, highly interconnected core containing three constituent communities sharing similar patterns of input connectivity. Intriguingly, several of the most important nodes in this connectome are little-studied neurons that are specialized for peptidergic neuromodulation. We anticipate that the *C. elegans* neuropeptidergic connectome will serve as a prototype to understand basic organizational principles of neuroendocrine signaling networks.

## Introduction

Understanding how behavior arises from neuronal interactions in the brain is one of the great challenges of modern neuroscience. In recent years, many projects have started to map the synaptic wiring diagrams (or synaptic connectomes) of simple and complex nervous systems to define their architecture at the cellular level^1–4^. Specifically, connectomics has aimed to identify all the individual units of the nervous system (i.e., its neurons) as well as all the connections between these units. In practice this involves reconstructing volumes of brain tissue using electron microscopy (EM), and then analyzing these images to trace neuronal processes and identify pre-and post-synaptic hallmarks of chemical synapses between identified neurons. Currently there are several endeavors of this type that are compiling complete connectomes for organisms important for neuroscience research, including *Drosophila*, *Platynereis*, *Ciona,* zebrafish, and mouse^2, 3, 5–7^.

Although most connectomics research focuses on synaptic connectivity, chemical synapses are only one of several paths through which neurons communicate in the brain. For example, many important interactions between neurons involve volume transmission, by which secreted molecules are released extrasynaptically and activate receptors on neurons that are not wired through synapses or gap junctions to the releasing neuron. Unlike synaptic and gap junction transmission, where signaling is restricted to neurons on either side of the synapse, volume transmission can mediate signaling across distances of microns^8–10^. Such extrasynaptic signaling has been described for classical neurotransmitters as well as monoamines, and is particularly prevalent for neuropeptides, which are released from dense core vesicles outside the synaptic active zone and act on a longer temporal and spatial scale^10, 11^. Although these extrasynaptic interactions are largely independent from the synaptic wiring, they play key roles in neural circuits and are thus critical for understanding the neural basis of behavior^12–18^.

Neuropeptides carry out particularly complex and important functions in the brains of all organisms^19–21^. Neuropeptides are thought to be among the most ancient neuronal signaling molecules, and their origin may in fact predate the actual evolution of neurons^22, 23^. Biologically-active peptides are usually formed from larger genetically-encoded precursors, which are enzymatically processed through proteolysis and/or chemical modification^24^. Although some neuropeptides have been shown to activate ion channels or transmembrane kinases, they typically bind to G protein-coupled receptors (GPCRs), which act through second messenger pathways to modulate diverse aspects of neuronal physiology^25^. Neuropeptide-activated GPCRs fall into two main classes, the rhodopsin-like (Class A) and secretin-like (Class B) families, and within these groups many receptors (e.g. oxytocin/vasopressin, neuropeptide Y/F, neuromedin U, somatostatin) show significant conservation across widely divergent animal phyla^26, 27^. Neuropeptide systems also play broadly conserved roles in the control of behavioral states, including those involved in feeding, sleep and arousal, reproductive behavior, and learning^14, 28–33^. In humans, the role of neuropeptides as neuromodulators has made this signaling system a highly sought-after target for new neuropsychiatric treatments. Currently 50 drugs acting via GPCRs have been approved by the FDA^34^, including several neuropeptide-GPCRs such as: orexin antagonists for treatment of insomnia^35^, a substance P antagonist for treatment of chemotherapy-induced nausea^36^. Besides new treatments are being developed like a GIP agonist as treatment for Alzheimer and Parkinson’s^37^. These promising therapies and the large number of remaining neuropeptide-GPCRs in humans^18, 38^ indicate the therapeutic potential of this molecular system.

The diversity and extent of neuropeptide signaling implies that the pathways for peptidergic communication can also be considered as a network. The genomes of all animals encode many, in some cases hundreds of neuropeptides, along with a similarly large number of GPCRs^18, 39–41^.

Moreover, in contrast to monoamines, which are typically expressed in only a small subset of neurons in the brain, neuropeptides are expressed extremely broadly; indeed, recent single-cell transcriptomic studies indicate that most if not all neurons in the mouse cerebral cortex express one or more neuropeptides as well as multiple neuropeptide-binding GPCRs^39, 42^. These data imply that peptidergic signaling underpins a dense and pervasive interaction network involving most if not all neurons of an animal’s nervous system^43, 44^. Since neuropeptide signaling is thought to be mostly extrasynaptic, the topology and structure of this wireless neuropeptide connectome is expected to be fundamentally distinct from that of the wired synaptic connectome^45^. However, in most organisms there is insufficient gene expression data mapped with single-cell resolution to identified neurons to map the structure of these extrasynaptic networks.

The nematode *C. elegans* represents an attractive animal to investigate the structure and organization of neuropeptide signaling networks due to its small and well-characterized nervous system. *C. elegans* was the first and is currently still the only adult organism with a completely mapped synaptic neuronal connectome, with each of its 302 neurons and approximately 2300 synaptic connections identified through electron microscopy (EM) reconstructions^46–48^. Despite its small size, the *C. elegans* nervous system has been found to share several structural features with that of larger animals. For example, the *C. elegans* connectome, like those of larger nervous systems, exhibits a small-world topology, with relatively high clustering paired with relatively short average path lengths^49, 50^. Likewise, the *C. elegans* nervous system is highly modular, with functionally segregated local clusters of high within-group connectivity^51–54^. Finally, the worm connectome contains a small number of highly connected hubs, which are interconnected in a core or rich club and facilitate communication between modules^55^; similar rich club topology has been observed in bigger brains, including the human cortex^56, 57^. Shared structural features are also apparent at the microcircuit level; for example, feed-forward motifs are significantly overrepresented in the nematode connectome, just as in the mammalian cortex^58–60^. Thus, insights gained from analysis of neuropeptide signaling networks in *C. elegans* may also reveal organizational principles that are conserved in larger brains.

Although the *C. elegans* nervous system is anatomically small, at the biochemical level its neuropeptide signaling pathways show similar complexity and remarkable conservation with humans and other larger-brained animals. Its genome contains 159 predicted neuropeptide precursor genes (NPP) (including 40 insulin-like peptides) that produce over 300 different neuropeptides^24, 61^.

Likewise, approximately 150 *C. elegans* genes encode GPCRs known or predicted to be activated by neuropeptides^25, 62^. These numbers are of a similar order of magnitude to the numbers of neuropeptides and peptide activated GPCRs in the human genome^18, 63^. Each *C. elegans* neuron not only expresses at least one neuropeptide and at least one neuropeptide receptor, as do vertebrate neurons,^18, 39, 40, 42, 43^ but each *C. elegans* neuron class expresses a unique combination of neuropeptide-encoding genes^40^, a notion that illustrates the tremendous potential complexity of neuropeptide signaling. Many *C. elegans* neuropeptides and cognate receptors are phylogenetically conserved across animal phyla and have clear human homologues, with some families (such as the RFamide peptides) having undergone expansion in the nematode lineage^64^. Thus, neuropeptide signaling in nematodes shows surprising conservation as well as similar diversity to neuropeptide signaling in the human brain, despite vast differences in neuron number and anatomical complexity.

Here we present a draft connectome of neuropeptidergic signaling in *C. elegans,* built by integrating gene expression, ligand-receptor interaction, and anatomical datasets. We identified predicted neuron-to-neuron signaling interactions mediated by 91 neuropeptide-receptor pairs, and by aggregating these individual neuropeptide-signaling networks generated a comprehensive network of peptidergic signaling in the *C. elegans* nervous system. This resulting neuropeptidergic connectome differs significantly in its structure from the previously-characterized wired connectome^47^; for example, it is denser, contains distinct hubs, high-weight edges, and autocrine signaling foci, and has a decentralized topology allowing direct communication among a large fraction of neurons. We expect this nematode neuropeptide connectome will serve as a prototype for understanding the organization of peptidergic signaling and how it interacts with the wired neural circuitry in other organisms, including those with much larger brains.

## Results

### Construction of a neuropeptidergic connectome

To generate a draft neuropeptide connectome, we integrated information from three datasets to infer potential pathways for neuropeptide signaling between individual *C. elegans* neurons (Figure 1A). To identify biologically relevant molecular interactions between individual neuropeptides and receptors, we used data from a large-scale reverse pharmacology screen which tested 55,386 potential neuropeptide-receptor interactions *in vitro* and identified 461 peptide-GPCR couples encoded by 55 neuropeptide and 57 GPCR genes^40^. To determine which neurons express each neuropeptide and receptor, we used publicly available single-neuron transcriptome data from the CeNGEN project, which described single-cell RNA-sequencing transcription profiles of all predicted neuropeptide and peptide-activated GPCR genes in *C. elegans*^40^. We validated these scRNA datasets with several CRISPR/Cas9-engineered reporter alleles of ligands and receptors. Finally, we used previously published EM anatomical reconstructions to assess physical constraints on potential diffusion of neuropeptides to their target receptors^48, 65, 66^. In constructing the neuropeptide network, we consider that two neurons (nodes) are connected by a directed edge if the first neuron expresses a particular neuropeptide ligand and the second expresses a paired receptor, subject to spatial constraints on signaling (Figure 1B).

**Figure 1.**
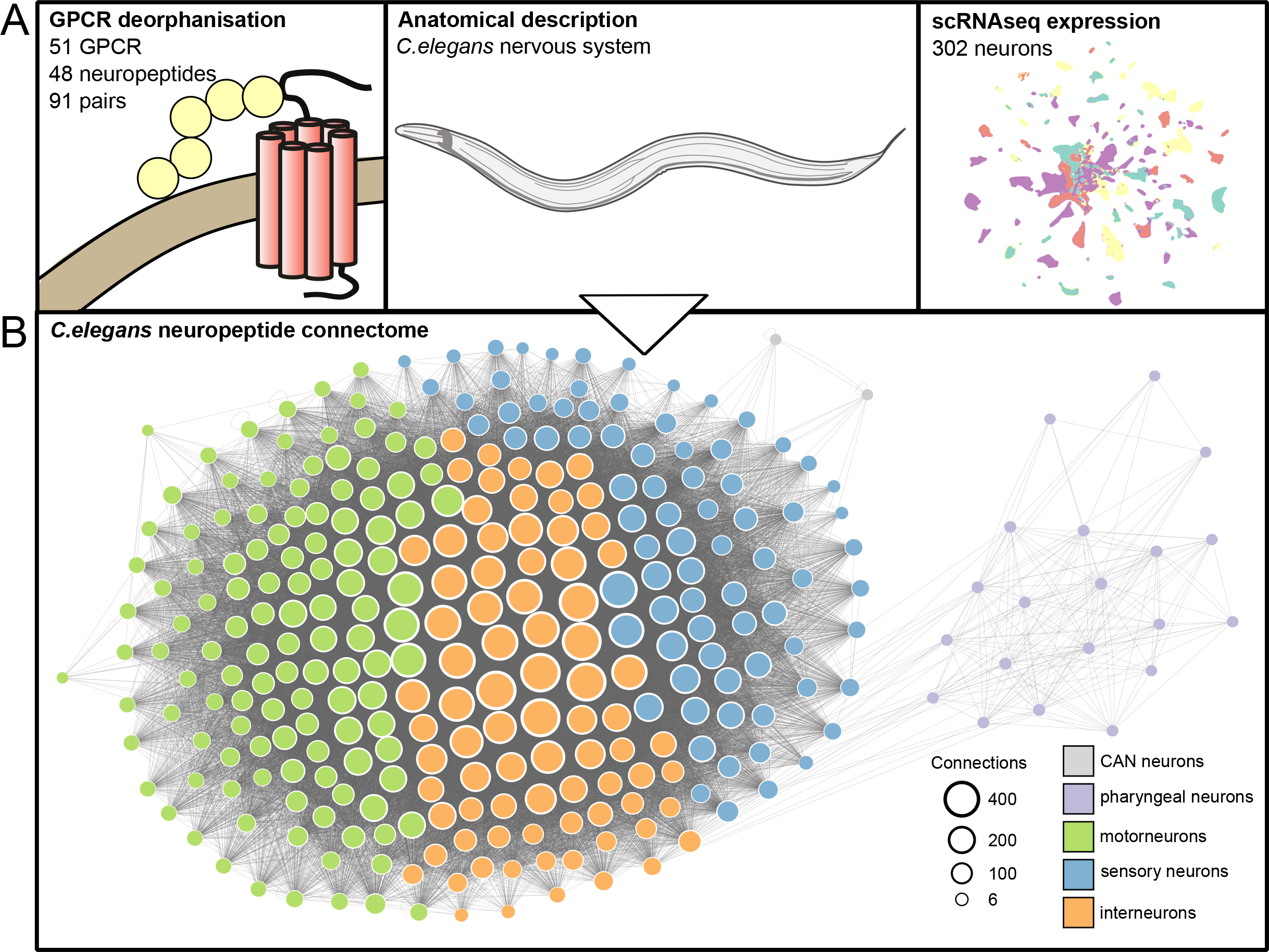
*In silico* reconstruction of the neuropeptidergic connectome of the *C. elegans* nervous system by combining *in vitro* neuropeptide-GPCR interaction, gene expression and anatomical datasets. (A) Datasets used to build the network *in silico.* Left: biochemical neuropeptide-GPCR interaction dataset defining 91 neuropeptide - G protein-coupled receptor (GPCR) pairs below 500 nM EC_50_ affinity, coupling 51 GPCRs to 48 neuropeptides^64^. Neuropeptides from the same precursor gene (NPP) are considered the same. Middle: anatomical and morphological description of the *C. elegans* hermaphrodite nervous system^48^. Right: single-cell RNAseq expression of the 302 *C. elegans* neurons, adapted from^40^. (B) Representation of the *C. elegans* nervous system connectome network in which all types of neurons (pharyngeal, interneuron, CAN neurons, motorneurons and sensory neurons) are connected through neuropeptidergic connections.

For each dataset, it was necessary to threshold for biologically relevant interactions.

For biochemical interactions, the raw dataset contained 461 neuropeptide-receptor pairs that showed dose-dependent activation with a half-maximal effective concentration (EC_50_) ranging between 0.1 pM to 22 μM^64^. At the gene level, these pairs were encoded by 55 neuropeptide and 56 GPCR genes, with 148 unique gene-gene interactions. In assessing neuropeptide-receptor pairings we initially opted for a conservative EC_50_ threshold of 500 nM, as many neuropeptide-receptor couples with EC_50_ values in this range have been validated *in vivo*^14, 25, 28, 32, 62^. By this criterion, we defined 91 individual neuropeptide-receptor gene couples, with a large number (51) of the predicted neuropeptide GPCRs having at least one identified ligand. The ligand-receptor interactions were complex, with several peptides activating more than one receptor (promiscuous receptors) and several receptors being activated by multiple peptides encoded by distinct precursor genes (versatile peptides)^64^.

We next sought to determine an appropriate threshold for the scRNA-based gene expression dataset. The transcriptomic data from the CeNGEN project has been differentially thresholded based on a ground-truth dataset of reliable gene expression from across the whole *C. elegans* nervous system using fosmid or receptor tagged reporters^40^. Each threshold was given a false discovery rate and a true positive rate depending on the number of correctly identified cells for a given gene by the scRNAseq analysis^40^. To determine the most appropriate threshold for our analysis, we obtained single-copy genomic knock-in reporters for 17 representative neuropeptide precursor genes and 9 representative genes for neuropeptide receptors and characterized their expression patterns comprehensively using the NeuroPAL marker strain (Figure 2A, B; Supplementary Figure 1). For both we found that threshold 4, although on some occasions overly stringent, was a good approximation of the expression pattern (Figure 2; Supplemental Table 1-3). Thus, although this stringent threshold has lower discovery power and thus may undercount the number of neurons expressing each gene, its stringency minimizes the likelihood that our resulting network will contain edges that do not represent authentic paths for potential neuropeptide signaling.

**Figure 2.**
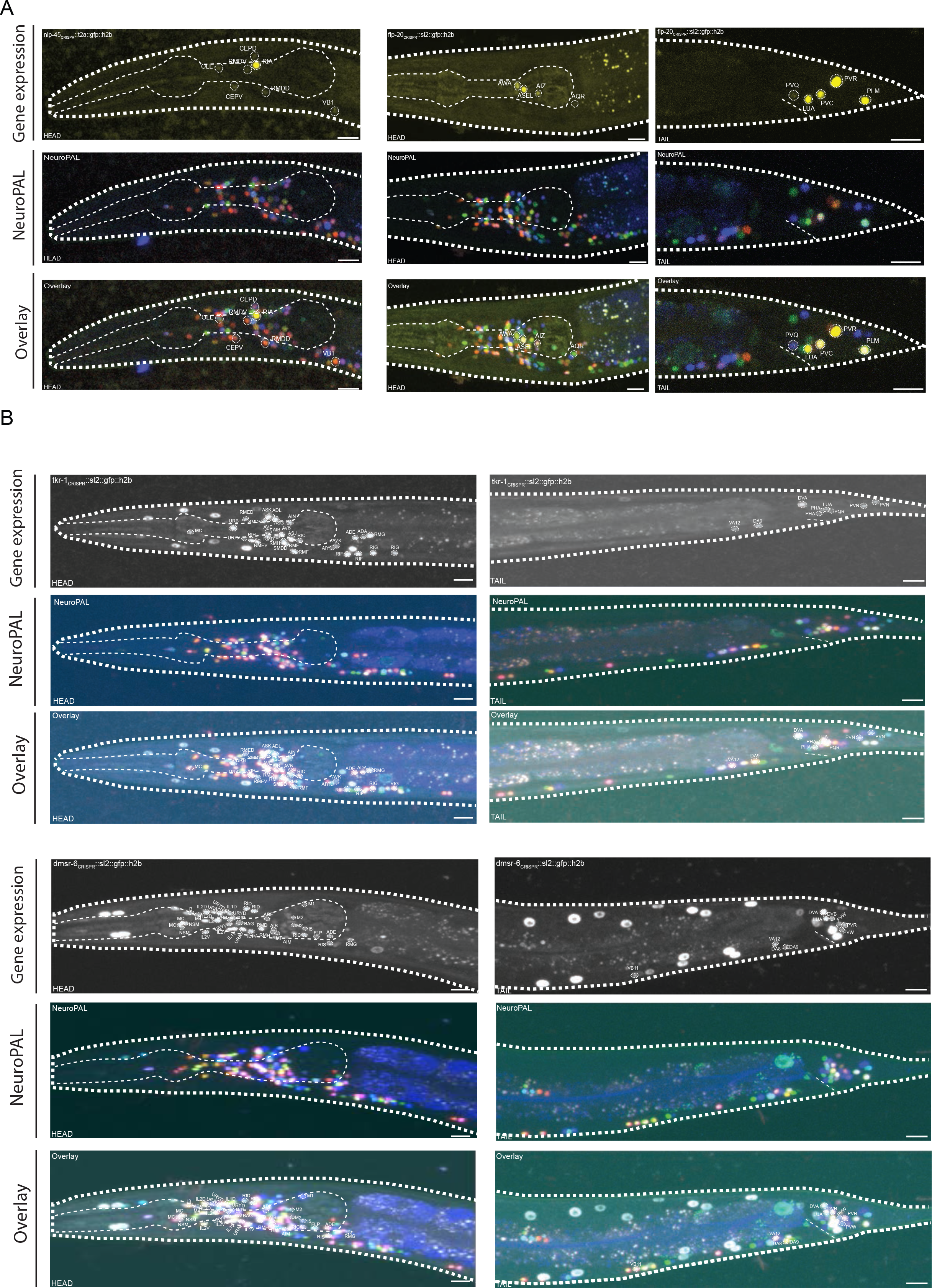
Single-copy genomic knock-in reporters corroborate the expression profiles of neuropeptide and GPCR genes obtained from scRNAseq. GFP-positive neurons were identified using the NeuroPAL multicolor transgene ^111^Figure pictures all segments (head, tail, vulva and midbody) showing neuronal expression, within which individual neurons are circled and labeled. Scale bars represent 10 µm. Tables of expression data and comparisons between cells identified in reporters and scRNAseq detection thresholds are in Supplementary Tables S1-3. (A) Fluorescent GFP reporters for the expression of two representative neuropeptide genes, *flp-20* and *nlp-45,* showing high correalation between reporter expression and CeNGEN scRNAseq expression. Data for additional neuropeptide gene reporters are shown in Supplementary Figure S1A. (B) Fluorescent GFP reporters for the two representative neuropeptide-activated GPCRs, expression of *tkr-1* and *dmsr-6*, showing high correalation between reporter expression and CeNGEN scRNAseq expression.^111^. Data for additional GPCR gene reporters are shown in Supplementary Figure S1B.

### Evaluating spatial constraints on neuropeptide signaling

These biochemical and gene expression datasets allowed us to infer which neurons express neuropeptide and receptor genes that could mediate a neuromodulatory interaction; however, the physics of diffusion and the neuroanatomy of the animal might potentially constrain some of these interactions *in vivo*. We therefore considered several possible models of the neuropeptidergic network based on different anatomical and spatial constraints (Figure 3B, C). We used EM reconstruction data for each individual neuron in the *C. elegans* nervous system to identify its anatomical location (Figure 3A, Supplementary Table 4) and the neuropil bundles into which their axons and dendrites project^48, 65, 66^. In the first and most permissive model, long-range signaling is permitted and neuropeptidergic connections can take place between any neurons in the system. In the second (“mid-range”) model, neuropeptidergic connections can occur only between neuronal processes in the same anatomical area, such as the head (including pharynx), tail and midbody (Figure 3A). In the third (“short-range”) model, we only considered potential peptidergic connections between neurons in the same neuronal process bundle, that is, between neurons whose processes overlap in the nerve ring, ventral or dorsal cord, or an auxiliary nerve. Finally, it has recently been shown that the nerve ring, into which more than half the neurons (168) project, can be divided into four layers or strata based on patterns of physical contact^67, 68^. We therefore also considered a fourth model in which neuropeptide signaling inside the nerve ring is constrained within individual strata and thus between neurons that make physical contact.

**Figure 3.**
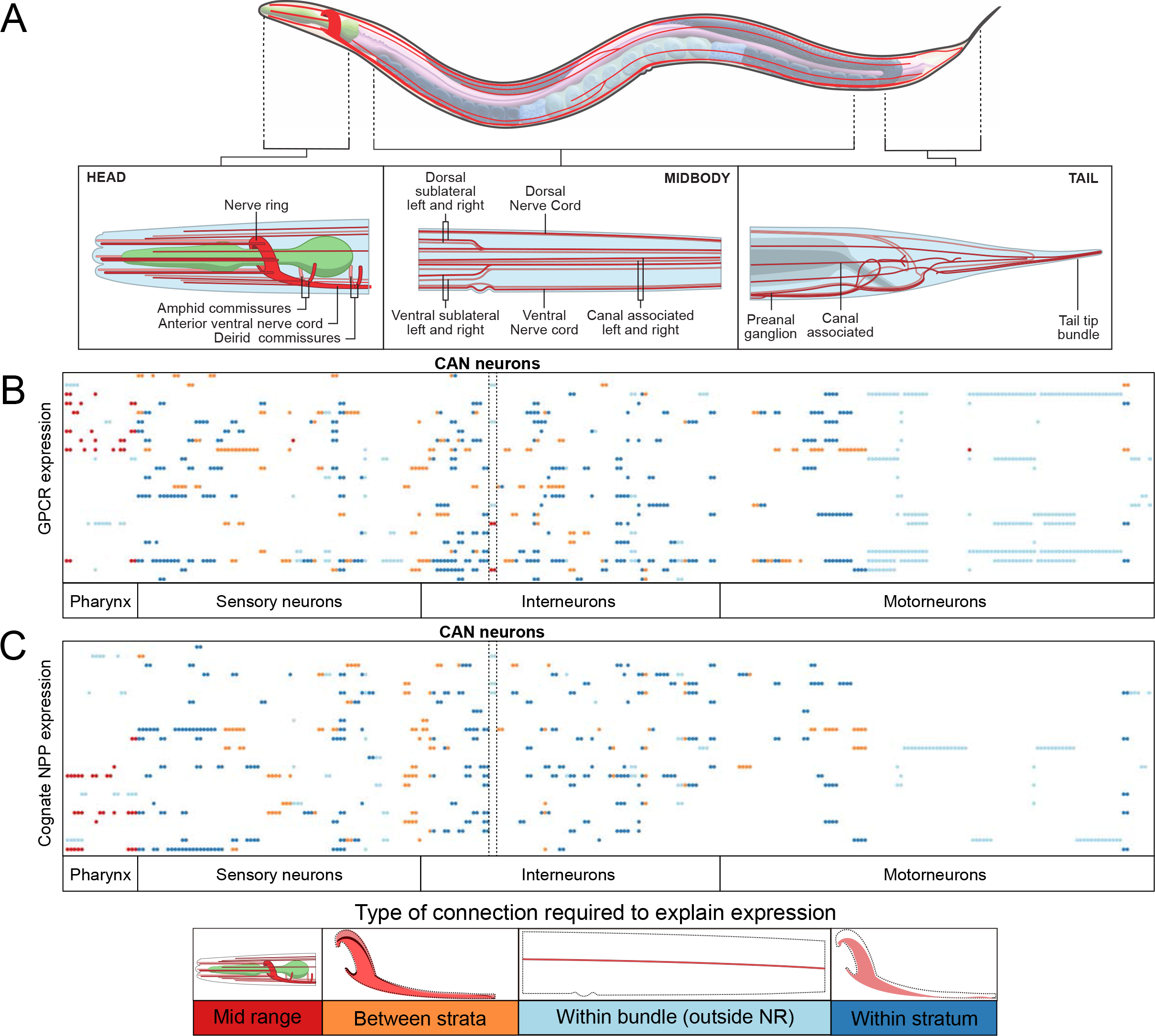
Analysis of neuropeptide and receptor expression allows assessment of the spatial scale of neuropeptide signaling. (A) Full anatomical description of the *C. elegans* hermaphrodite nervous system^112^. Neuronal bundles represented in red, and the pharynx in green. Short-range connections are defined as occurring within the same neuronal bundle (axons and dendrites considered equally). Mid-range connections occur between neurons whose processes are present in one of following distinct anatomical areas: head (ranging from nose to deirid commissures including the ventral and retrovesicular ganglion), midbody (between but not including deirid commissures and the preanal ganglion), or tail (from and including the preanal ganglion to the tail’s tip). Classifications of individual neurons are in Supplementary Table S4. (B) Expression matrix for 23 GPCRs activated *in vitro* by a single neuropeptide ligand. Neurons are sorted by neuron type on the x axis and the expression of each GPCR on the y axis. Colors indicate the range of diffusion distance required for communication with at least one ligand-expressing neuron: blue indicates that the receptor expressing neuron makes contact with a neuron expressing its ligand, either within the same stratum of the nerve ring (dark blue) or in a thinner neuronal bundle (light blue); orange indicates short-range connections between strata in the nerve ring (i.e. between neurons in same bundle but not in physical contact); red indicates connections between neurons in the same anatomical area but not in the same bundle. Nerve ring stratum as defined^67, 68^ (C) Expression matrix for 23 neuropeptide precursor genes with a single target receptor in vitro. Neurons are sorted by neuron type on the x axis and the expression of each neuropeptide gene on the y axis. Colors indicate the range of diffusion distance required for communication with a receptor-expressing neuron, as described above.

To evaluate each model, we investigated whether the expression patterns of receptors and ligands were consistent with neuropeptide signaling being restricted as described above (Figure 3B, C). For example, if neuropeptide signaling occurred only between neurons in physical contact (as in model 4) we would expect neurons expressing a given receptor to project to the same nerve ring strata as neurons expressing its ligand. To simplify this analysis, we focused on the neuropeptide-receptor couples in which the receptor has only one ligand and the neuropeptide only binds to that receptor, assessing whether, under the constraints of each model, there would be receptor-expressing neurons that were unable to receive a signal from any neurons expressing its ligand. Conversely, we also analyzed couples where the ligand activated only one receptor, asking whether under a given model there were ligand-expressing neurons that could not communicate with cells expressing its receptor.

This analysis appeared to argue against the most restrictive model, in which ligands could not diffuse between nerve ring strata. Specifically, we observed many examples in which a neuropeptide receptor was expressed only in strata that did not express its specific ligand (Figure 3B, C, orange points). For instance, while the *capa-1* neuropeptide precursor gene is expressed only in the ASG sensory neuron, which projects its axon into the fourth nerve ring stratum, the gene for its receptor NMUR-1 is expressed in all four strata^32, 69^(Figure 3B, C). Thus, NMUR-1 receptors in strata 1-3 must be activated by CAPA-1 peptides that diffuse from their release site in stratum 4. Overall, 37 of the 41 receptors analyzed were expressed in at least one neuron making no contact with a neuron expressing its ligand. Likewise, 32 of the neuropeptide precursor genes were expressed in at least one neuron making no contact with neurons expressing its receptor (Figure 3C, yellow points). Since the nerve ring strata are not separated by glial or other barriers, and experimental evidence^14, 32^ indicates neuropeptides can indeed travel between strata, it is reasonable to infer that neuropeptides are not likely to be constrained by anatomical layers in the nerve ring or other neuronal bundles.

We likewise observed cases in which the expression of a neuropeptide or receptor gene implies mid-range signaling between distinct nerve bundles in the same body region (Figure 3B, C). For example, the *frpr-7* receptor gene is expressed in multiple pharyngeal neurons, while its ligand FLP-1 is released exclusively from the AVK neuron whose processes lie in the nerve ring and ventral cord^13^. Thus, FRPR-7 receptors in the pharynx must be activated by peptides released by AVK in the nerve ring, consistent with published evidence that neuropeptides can signal between head neurons and pharyngeal neurons^13, 70^. Indeed, for a majority (14/22) of the receptors expressed in pharyngeal neurons, their ligand was expressed only outside the pharynx (Figure 3B, red points). Likewise, receptors such as NPR-3 and TRHR-1 are expressed in the CAN neurons, yet their ligands FLP-15 and NLP-54 respectively are not expressed in any neurons projecting into the canal-associated nerve^71^ (Figure 3B, C, CAN neurons labelled). Thus, we hypothesize that at least some neuropeptide signaling occurs between different nerve bundles, in particular between the pharynx and the nerve ring and between CAN and other nerves in the body. We therefore focused our subsequent analysis on model 2, consisting only of stringently selected short-range interactions, and model 3, which includes mid-range signaling between nerve bundles in the same body region.

### Neuropeptide networks exhibit diverse topologies

Based on these criteria, we first constructed network graphs between neuropeptide-expressing and receptor-expressing neurons for each of the 91 individual neuropeptide/receptor couples in our dataset (Supplementary Figure 2-3). These networks were filtered by removing edges between neurons that did not project an axon or dendrite into the same process bundle (for the short-range network) or into the same body region (for the mid-range network). In their short-range version, 78 of these ligand-receptor couples formed a single connected network, whereas 13 formed networks with two or even three disconnected components (Figure 4B, Supplementary Figure 2). In their mid-range versions, all 91 couples formed a single connected network (Supplementary Figure 3).

**Figure 4.**
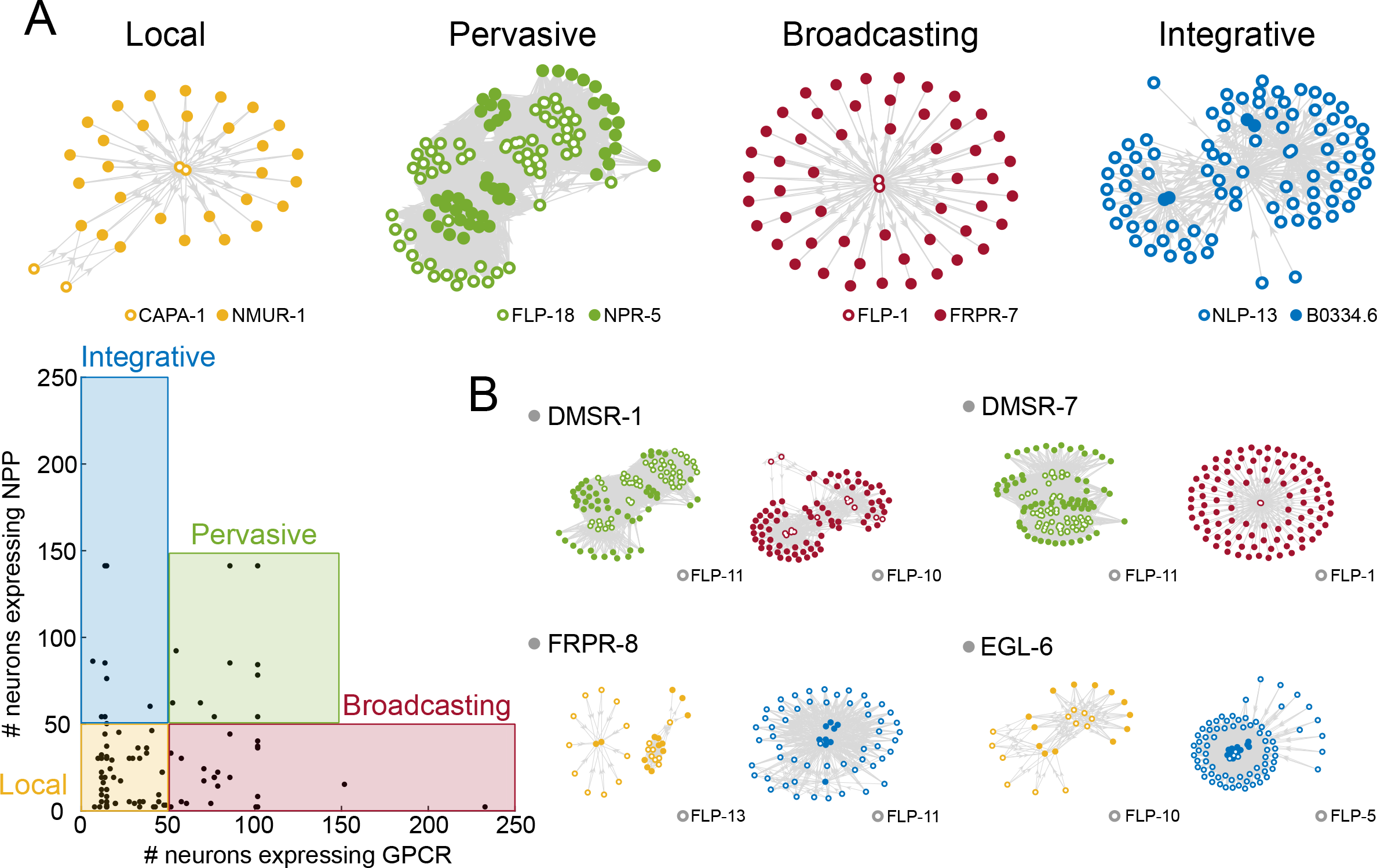
Assortativity analysis of 91 individual NPP-GPCR networks shows different topologies depending on the NPP-GPCR interaction. (A) Representation of network topologies depending on assortativity, reflecting the preference of neuropeptides to signal to GPCRs that are expressed in the same number of neurons. Networks are classified either as local, pervasive, broadcaster or integrative. Bottom left: scatter plot showing the number of neurons expressing a particular GPCR versus the number of neurons expressing the corresponding NPP gene for each of the 91 individual networks. Local networks show restricted NPP and GPCR expression in 50 neurons or less. Pervasive networks have broad NPP and GPCR expression in more than 50 neurons. Broadcaster networks show broad GPCR (˃50 neurons) but restricted NPP expression (≤50 neurons), while integrative networks display broad NPP (˃50 neurons) and restricted GPCR expression (≤50 neurons). Filled circles indicate receptor presence in that neuron and empty circles indicate neuropeptide release from that neuron. Left top: example of local network for CAPA-1/NMUR-1 that has been linked to learning^32^. Left middle top: example of pervasive network FLP-18/NPR-5 of which the receptor is implicated within locomotory behavior^119^. Right middle top: example of broadcaster network FLP-1/FRPR-7 that modulates the interaction between food sensation and locomotion^13^. Right top: example of integrative network NLP-47/GNRR-1, the receptor of which is linked to egg-laying behavior ^120^ and is only activated by NLP-47 *in vitro* ^64, 120^. Diagrams of all 91 networks are in Supplementary Figures S2 and S3. (B) The topology of the network is defined by the individual NPP-GPCR pairing. Networks of the same receptor can show different topologies depending on its neuropeptide ligand. In each example, the same receptor can form both pervasive and broadcasting networks (DMSR-2, DMSR-7) or both local and integrative networks (FRPR-8 and EGL-6) depending on the activating ligand.

We observed a diverse range of topologies in the networks for individual peptide-receptor couples. One topological measure in which the different networks varied was their assortativity, or the extent to which nodes are preferentially connected to nodes of similar degree. Using the conservative short-range model, we observed that most of the networks, including many of those highly conserved in evolution, were local networks, in which both ligand and receptor were expressed in relatively restricted sets of neurons (Figure 4A, Supplementary Figure 2). The neurons in these networks showed relatively low average degree and often encompassed only a subregion of the total nervous system. In addition, we found 8 highly disassortative *integrative* networks, with many low-degree peptide-releasing neurons signaling to relatively few high-degree receptor-expressing neurons (Figure 4A). We also observed 23 disassortative *broadcasting* networks that are characterized by a small number of high-degree peptide-releasing neurons that signal to many low-degree receiving neurons with broadly expressed receptors (Figure 4A, Supplementary Figure 2). Interestingly, both integrative and broadcasting networks involved promiscuous receptors with multiple ligands, specifically FRPR-8, EGL-6, DMSR-7, and DMSR-1. DMSR-7 and DMSR-1 also figured prominently in a fourth category of more assortative, pervasive networks, in which both the ligand and receptor show broad expression and most neurons thus show relatively high degree (Figure 4A, B). Relaxing the spatial restrictions on our model had relatively modest impact on network topology, with four networks that are local in the short-range model becoming either integrative (one network) or broadcasting (3 networks) in the mid-range model (Supplementary Figure 3). Thus, the topologies of neuropeptide networks appear relatively robust to our assumptions about the spatial scope of neuropeptidergic signaling.

### The neuropeptide connectome is a decentralized, dense network

By aggregating the networks from the individual neuropeptide-receptor couples, we next compiled complete neuropeptide connectome networks based on short-range or mid-range signaling (Figure 5). Even for the more conservative short-range network, in which signaling was restricted to neurons with processes overlapping in the same bundle, the density of connections was remarkably high, with more than a third of all possible connections or edges (0.3437) present in the network.

**Figure 5.**
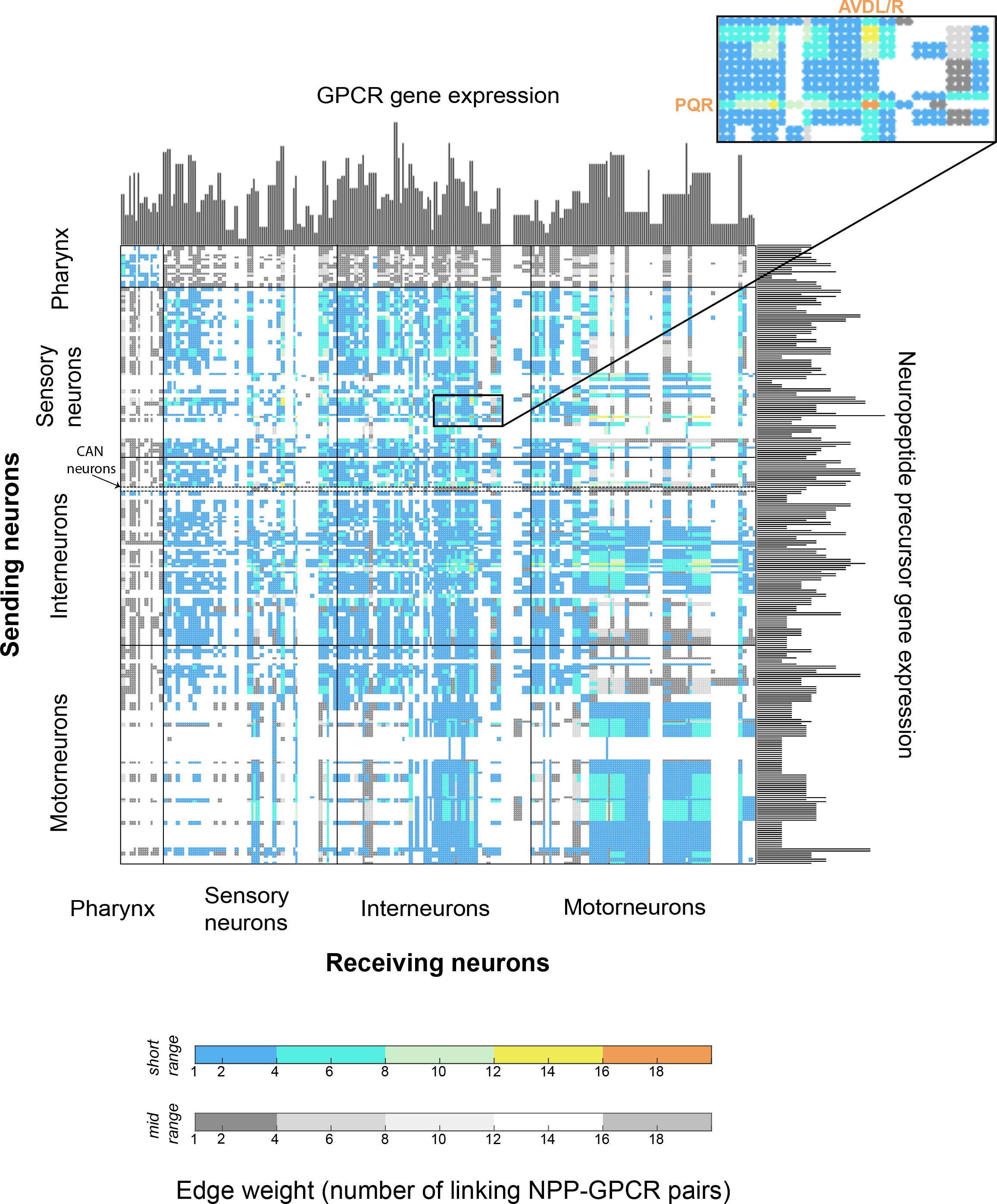
The aggregate neuropeptide connectome in *C. elegans* connects all 302 neurons in a dense network. Adjacency matrix representation of the aggregate 91 NPP-GPCR pair network considering short-range (color) and mid-range (gray) diffusion models. Axes represent the number of NPP and GPCR genes per neuron. Virtually all neurons express at least 1 NPP and 1 GPCR gene, the only exception of which is the IL1 neuron class that does not show GPCR expression. Neurons can communicate with each other using up to 18 different NPP-GPCR pathways, a feature conserved in both short-and mid-range diffusion models. Dense and diverse connections are mainly seen from oxygen sensing neurons to inter-and motorneurons. 5% of all connections are putative autocrine connections in which a single neuron co-expresses both NPP and GPCR genes of a cognate pair. Supplementary Figure 4 shows adjacency matrices for long-range (A) and mid-range (B) models.

Allowing mid-range connections (0.4429) or long-range connections (0.5929) increased this density further (Supplementary Figure 4A, B). By comparison, the *C. elegans* synaptic (0.0251) and monoamine (0.0236) networks were far less dense, fourteen-fold lower than the short-range neuropeptide network. We also computed edge weights for the connections in the neuropeptide networks based to the number of neuropeptide-receptor pairs capable of signaling between two nodes. We observed that a large number (35% in the short-range network) of neuron pairs were connected by only a single neuropeptide-receptor interaction, but some (9%) were connected by 6 or more different peptide-receptor couples (Figure 5). In the most extreme case, we observed that two neurons, the AVD premotor interneurons and the PQR oxygen-sensing neurons, were linked by 18 different neuropeptide-receptor pairs, suggesting extraordinarily complex patterns of signaling could exist between these cells (Figure 5, highlighted). Interestingly many other high-weight, biochemically complex connections occurred between neurons of the oxygen sensing circuit and were mediated by a common set of promiscuous receptors (DMSR-1, DMSR-7 and FRPR-8) and versatile neuropeptides (FLP-4, FLP-9, FLP-10 and FLP-13). These neurons are thus connected by multiple neuropeptidergic channels of communication, potentially allowing complex regulation by context and experience.

A striking feature of the midrange network in particular was its integration of neurons disconnected from the synaptic connectome. For example, the 14 neuron classes of the pharyngeal nervous system form a heavily synaptically interconnected, functionally autonomous network akin to vertebrate enteric nervous systems, which is topologically isolated from the rest of the nervous system^65, 72, 73^. We find that unlike the wired connectome, there are no neuropeptidergic networks that define circuitry exclusively in the pharynx; to the contrary, pharyngeal neurons are fully integrated into the somatic nervous system via strong reciprocal interconnectivity (Figure 5). All pharyngeal neuron classes receive 90% or more of their incoming connections from outside the pharynx. Some classes of pharyngeal neurons also broadcast extensively to the somatic nervous system, with several (I1, I3, I4, I5, M5 and NSM) having more than 100 outgoing connections (more than 90% of their total) to non-pharyngeal neurons. Likewise, it is notable that the CAN neurons, which completely lack chemical synapses, show strong and reciprocal neuropeptidergic connectivity with the rest of the nervous system, indicating that this unusual neuron class is well embedded in the neural network (Figure 5, CAN highlighted).

We next investigated topological features of the aggregate neuropeptide network. In particular, we analyzed the degree nodes of the network, defined as the number of incoming (in-degree) and outgoing (out-degree) connections made by each neuronal node. Degree is often an indicator of functional significance in networks, since high-degree nodes or hubs often play key functional roles in the brain and other complex systems. As expected from the high density of neuropeptide signaling, the average degree of both the short-range (Figure 6A) and mid-range networks (Figure 6B) were significantly higher than for the previously characterized synaptic (Figure 6C) and monoamine (Figure 6D) networks. Moreover, when we analyzed the distribution of degree across the neuropeptide networks, we observed relatively flat curves with many high-degree nodes; more than half the neurons in the short-range network and nearly two thirds in the mid-range network had a degree higher than 200, indicating that they have in and out connections with more than a third of all other neurons (Figure 6A, B). In contrast, the synaptic network was much more centralized, with only 10 neurons having degree greater than 50 (Figure 6C); similarly, the monoamine network had only 18 high (>50) degree neurons (Figure 6D). The best-connected neurons in the neuropeptide network have exceptionally high in-degree as well as out-degree, indicating that their status as hubs depends as much on incoming connections as outgoing connections, and as much on their expression of broadly-signaling neuropeptides as on integrating GPCRs (Figure 6A, B); this contrasts with the monoamine network, where the hubs are exclusively monoamine-releasing neurons with a high out-degree (Figure 6D).

**Figure 6.**
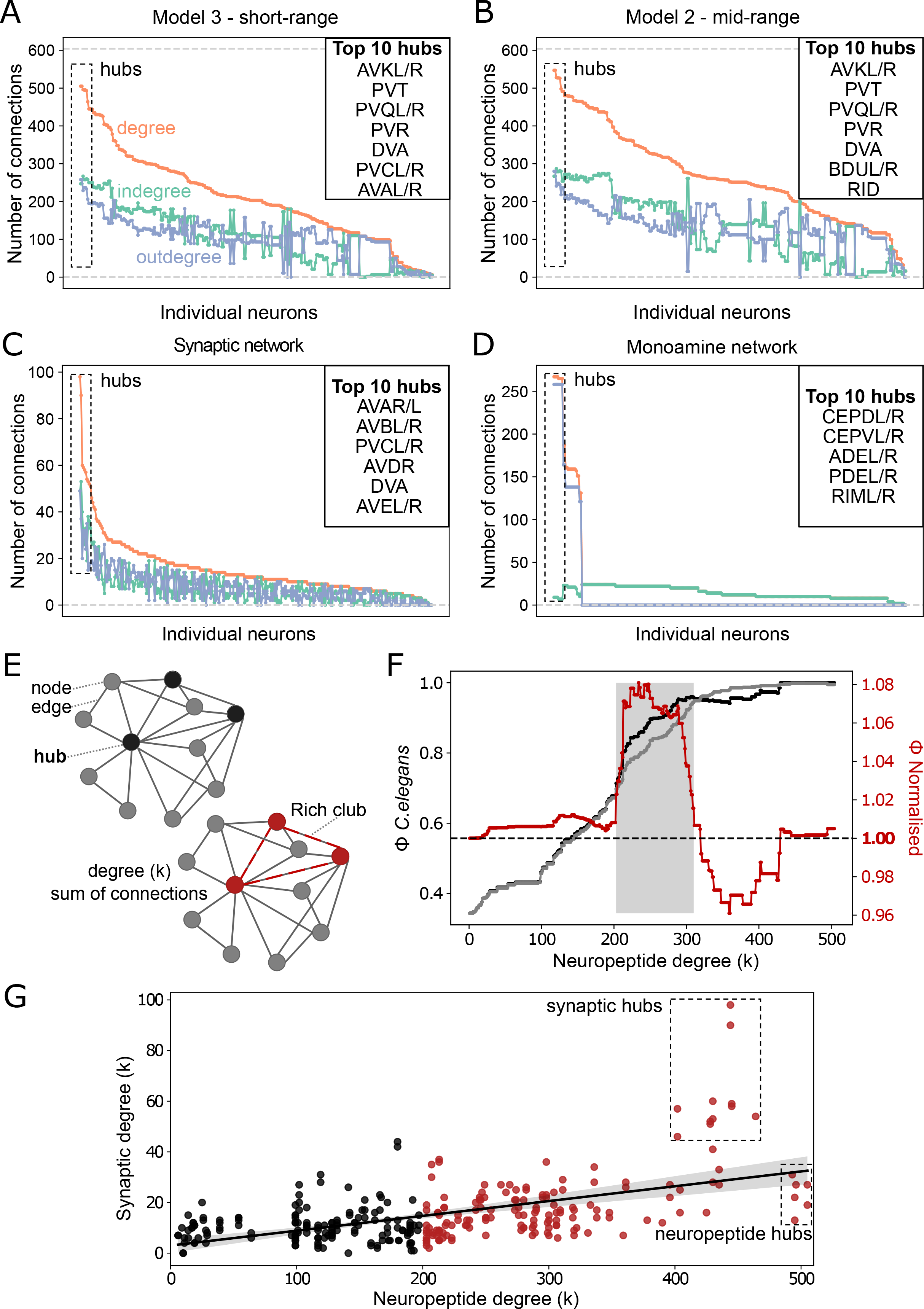
Topological analysis of the *C. elegans* nervous system highlights neuropeptide signaling hubs and a large rich club. (A) Degree distribution of the short-range model aggregate neuropeptide network. Degree (sum of incoming and outgoing connections) is shown in orange, the indegree (sum of incoming connections) is shown in green and the outdegree (sum of outgoing connections) is shown in purple. . The 10 hubs with the highest degree are indicated, including hubs of the synaptic connectome (AVAL/R, PVCL/R and DVA) as well as 6 neurons with specialised roles in the neuropeptide signaling network (AVKL/R, PVT, PVQL/R, PVR). (B) Degree distribution of the mid-range model aggregate neuropeptide network. Similar to the short-range model, degree distribution is equally influenced by both in- and outdegree: the 10 hub neurons of highest degree are indicated. (C) Degree distribution of the synaptic network of *C. elegans*. Compared to the neuropeptide connectome, the synaptic connectome contains a much smaller number of high-degree hub nodes. (D) Degree distribution of the monoamine network of *C. elegans.* The overall degree distribution shows two distinct phases, correlating with the network having a star-like topology centered around a small core of high degree hubs. This topological feature is further reflected in the low outdegree of most neurons. (E) Conceptual network representation highlighting: node (neuron), edge (connection between neurons), degree (k, sum of incoming and outgoing connections), hub (highly connected neurons), and rich club (group of hubs that connect more to one another than to the rest of the network). (F) Rich club of the *C. elegans* neuropeptidergic connectome. The rich club coefficient Φ(k) for the *C. elegans* neuropeptidergic network is shown in black and the randomized rich club curve <λ_random_(k) generated by averaging the rich club coefficients of 100 randomized versions of the *C. elegans* neuropeptidergic network that preserve degree distribution) is depicted in gray. The red curve is the normalized coefficient. <λ(k) ý <λ_random_(k) + 10σ over the range 203 :: k :: 298, indicating the rich club. The large number of connections in the range k > 298 make the random networks more unreliable but since Φ(k) is very close and mostly higher than <λ_random_(k) + 10σ we can infer that all neurons where 203 :: k are part of the rich club. This means the short-range aggregate neuropeptide network has a rich club of 156 neurons (166 in the mid-range, Suppl. Figure 6) (G) Correlation between synaptic and neuropeptidergic degrees. A positive correlation was observed over all neurons (r = 0.54, p = 3.1 e^-14^). Dark red dots indicate neurons in the *C. elegans* neuropeptidergic rich club. Synaptic hubs, which also exhibit a high neuropeptide degree (k>400) (ranking among the top 30 neuropeptide hubs) are highlighted. Also highlighet are 6 neuropeptidergic hubs, with low synaptic degree (k < 30, below the correlation line), as well as low monoamine (22 > k 2 10) and gap junction (14 > k 2 3) degrees (Suppl. Figure 6), appear to be specialized for neuropeptide signaling.

### The neuropeptide connectome contains a highly connected core with unique peptidergic hubs

A significant feature of many networks is the so-called rich club property. In such networks, the most highly connected hubs form more connections between themselves than expected based on their high degree alone; these hubs therefore comprise a rich club or core of the network that facilitates communication between more peripheral network nodes. The *C. elegans* synaptic connectome, for example, contains a rich club consisting of 11 premotor interneurons that appear to play important roles in driving global brain states^55, 74^. Remarkably, we found that the neuropeptide connectome also shows the rich club property, but its rich club consists of 156 neurons (166 in mid-range network), more than half of the nervous system (Figure 6F). Within the rich club the density of connections is 0.6834 (p<0.00001), more than double the density of the overall network (0.3427).

Interestingly, compared to the wired synaptic connectome, the neuropeptide connectome as a whole has significantly higher clustering and reciprocity (i.e. neurons are more likely to connect in both directions) but lower modularity and disassortativity. This finding further supports the notion of a decentralized neuropeptide connectome, with a broad-based densely-connected core allowing direct communication between a large fraction of neuron pairs.

To further investigate the relationship between neuropeptide and synaptic signaling, we looked at the correlation between synaptic and neuropeptidergic degree (Figure 6G). We observed that neuropeptide degree and synaptic degree were positively correlated (p < 0.0001, r = 0.53). We further observed that neurons with a high synaptic degree were among the most important neuropeptidergic hubs; for example, all 11 members of the synaptic rich club (DVA, PVCL/R, AVAL/R, AVBL/R, AVDL/R, AVEL/R) were among the 25 highest-degree nodes in the short-range neuropeptide network (Figure 6G). However, there were also several neurons of very high degree in the neuropeptide connectome but unexceptional synaptic degree (Figure 6G); these neurons can be described as specialized for neuropeptide signaling. Six neurons (AVKL/R, PVQL/R, PVT and PVR) had higher short-range neuropeptide degree than any of the synaptic rich club neurons (Figure 6A); if mid-range signaling is considered these same six neurons remain the highest-degree hubs (Figure 6B). Interestingly, the AVKL/R and PVT neurons are notable for expressing no classical neurotransmitters or monoamines^75^, while the PVQL/R neurons appear anatomically specialized for neuropeptide signaling due to a preponderance of dense core vesicles^14, 76^. AVKL/R, and PVT have been previously linked to the control of global behavioral states related to sleep and arousal^14, 73^, but the functions of the other neuropeptide hub neurons PVQ and PVR are not well-characterized.

### The neuropeptide connectome core has a defined mesoscale structure

To further probe the structure of the neuropeptide connectome we investigated whether the network contained modules or other forms of mesoscale substructure. We first applied standard methods for modular decomposition, but the high density of the network precluded the identification of any discrete modules, unless we aggressively filtered out lower-weight edges where neurons were connected only by a small number of neuropeptide-receptor interactions. However, we hypothesized that other types of meso-scale structure may be present^52, 77^. We wondered for example whether we could identify subgroups of neurons with similar patterns of incoming and outgoing neuropeptide connections. We therefore applied dimensionality reduction methods (Principal Component Analysis (PCA) and t-SNE) to the connectivity matrix of the aggregate neuropeptide networks to identify groups of neurons that share a connectivity pattern. This analysis highlighted 3 clearly defined groups of neurons that receive similar incoming connections, along with a more diffuse cloud of neurons with more variable connectivity (Figure 7A, B). Interestingly the three clusters comprise most neurons of the neuropeptidergic network (155 of 302), 76% of which are also members of the neuropeptidergic rich club (112; Figure 7A, B). Interestingly, each group receives a characteristic range of incoming connections, and this alone is enough to significantly sort them into these groups (p<0.0001) (Figure 7D): group 1 neurons have indegree of 134-156 (mean rank 162.2), group 2 neurons have an indegree of 252-282 (mean rank 276.5), group 3 neurons have an indegree of 169-212 (mean rank 222.9), and neurons in the unclustered cloud have an indegree below 169 (mean rank 77) (Figure 7D).

**Figure 7.**
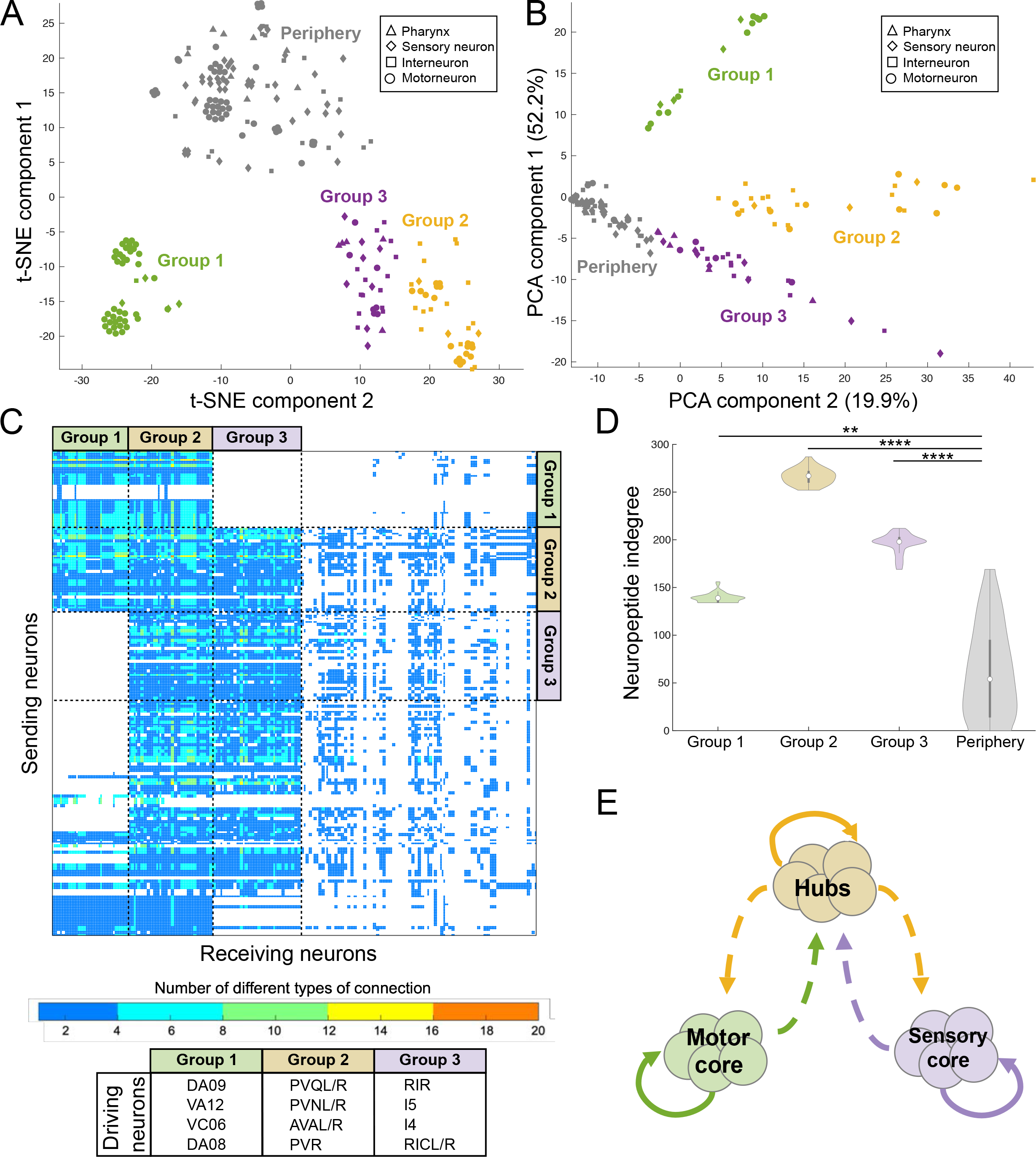
Mesoscale structure of the neuropeptide connectome (A) t-SNE dimensionality reduction of the adjacency matrix of the mid-range aggregate network (Euclidean distance, perplexity 30). This plot we identifies 3 clear clusters (1, 2, 3) which encompass 112 of out the 166 mid-range neuropeptide rich club neurons), in addition to a periphery of loosely clustered neurons. Datapoint markers represent their neuronal classification. (B) PCA dimensionality reduction of the adjacency matrix of the mid-range aggregate network (Euclidean distance, perplexity 30). Neurons in this plot are colored based on that of the groups defined in (A). A clustering pattern similar to that following t-SNE appears, highlighting the robustness of these defined groups across dimensionality reduction techniques. Datapoint markers represent their neuronal classification. (C) Mid-range aggregate network sorted in both dimensions (sending and receiving neurons) based on neuronal clusters defined in (A) and (B). Neuronal clusters divisions are highlighted with discontinuous lines. Group 3 does not receive nor send connections to group 1. All three groups, specially 2 and 3, send a large number of connections to peripheral neurons while not receiving significant input from the periphery. Group 2 is the only group that sends and receives connections to and from all 4 neuronal clusters. The neurons with higher weight of connections (number of NPP-GPCR pairs used to make the connection) are mainly found in groups 2 and 3. The neurons defining the patterns that drive the clustering of each group are shown at the bottom: group 1 is mainly driven by the connections of DA, VC and VA terminal motorneurons; group 2 is mainly driven by the connections of neuropeptidergic hubs PVQL/R and PVR, synaptic hubs AVAL/R and PVNL/R neurons; group 3 is mainly driven by the connections of pharyngeal neurons I5 and I4 and RIR, RICL/R interneurons. (D) Violin plots showing indegree values for the 3 clusters and the periphery. Indegree cleanly defines the 3 groups and the periphery (median group 1: 139, median group 2: 267, median group 3: 198, median periphery: 54), and therefore is the determinant factor in the definition of these groups. Significances determined by Kruskal-Wallis test followed by Tukey-Kramer test for multiple comparisons with rank sums. The mean ranks for the 4 groups were shown to be significantly different (mean rank group 1: 162, mean rank group 2: 276, mean rank group 3: 223, mean rank periphery: 77). n.s. not significant; \*\*\**p* ≤ 0.001. (E) Diagram showing the organization of connections between the 3 defined clusters. Group 1 contains the main motorneurons core. These connect between themselves, with the periphery and with group 2. Group 2 contains the main interneurons core including the top 10 hubs. These neurons connect to almost every other neuron in the nervous system. Group 3 contains the main sensory neurons and pharyngeal neurons core. These neurons connect to group 2 neurons and the periphery but do not connect to group 1 neurons. This structure indicates an intrinsic organization of the *C. elegans* neuropeptidergic connectome rich club and how it coordinates the function of the overall neuropeptidergic connectome.

These groups also diverge in the neuron types that form them and to which they are connected (Figure 7C, E). Group 1 is mostly motorneurons (motor core), including those involved in locomotion (VA, DA, VB, and DB)^78^ and the grouping of these neurons is driven by inputs from the posterior touch mechanosensory neurons PVM and PLML/R^79^ and the interneuron PVWL/R (Figure 7C).

Group 2 neurons are a mix of interneurons and motorneurons, including all top neuropeptidergic hubs (hubs core), that receive connections from every other neuron type, with the neuropeptidergic hubs (PQR, PVT and PVR) particularly important drivers for the grouping (Figure 7C). Finally, group 3 neurons are a mix of pharyngeal neurons, interneurons, motorneurons and sensory neurons (sensory core) that receive connections from every other neuron type but motorneurons, with RIR and pharyngeal neurons I5 and I4 driving the cluster (Figure 7C). Interestingly, although the characteristic neural inputs of the groups differed substantially, the most important (highest indegree) neuropeptide-receptor interactions for each group, involved versatile neuropeptides FLP-9, FLP-11 and FLP-16 and promiscuous receptors DMSR-7 and DMSR-1 (Figure 7C), leading to interconnections between groups (Figure 7E). For example, Group 1 forms connections with itself, Group 2 and with the unclustered cloud but not with Group 3; Group 3 forms connections with Group 2 and unclustered cloud neurons but not Group 1, and Group 2 forms connections with all other groups. Thus, Group 2 serves as a link with Groups 1 and 3, which have few direct connections with each other.

### Co-expression of neuropeptides with their target receptors generates autocrine pathways

Signaling cascades, in which a neuropeptide receptor is specifically co-expressed with a non-cognate peptide whose release it controls, are a classic hallmark of neuroendocrine pathways. To investigate whether the *C. elegans* neuropeptide connectome contains such cascades, we evaluated co-expression between neuropeptide and receptor genes. We performed a Fisher’s test to look for neuropeptide genes and GPCR genes that are co-expressed in the *C. elegans* nervous system more often than expected from their individual expression frequencies. We identified 121 peptide-receptor pairs meeting this criterion (Supplementary Figure 5), only 5% of which corresponded to cognate neuropeptide-receptor pairs (autocrine connections). Using these peptide-receptor pairs as nodes and the neuropeptide-receptor interaction data to form edges between them, we built a network of overrepresented signaling cascades within the larger neuropeptide connectome (Supplementary Figure 5B). This network is fully connected and provides a simplified view of how neuropeptide signaling pathways interact within the nervous system.

Co-expression of a neuropeptide receptor with its own ligand will generate a self-loop or autocrine connection, in which release of the peptide can signal back onto the sending cell. If we consider all cases where a peptide and its receptor are co-expressed in a given neuron, we find that 58% of the *C. elegans* nervous system harbors putative autocrine peptide connections (Figure 8A and 8B). Autocrine signals appear most prevalent in neurons categorized as either interneuron or motorneuron (Figure 8B), although two of the three types with the largest number of autocrine connections (URX and PQR) are sensory neurons. Promiscuous receptors participate more in autocrine peptide signaling (Figure 8C, Suppl. Figure S6), the most prominent of which are DMSR-1 and DMSR-7 (Figure 8C). We observed a strong positive correlation between the number of different autocrine peptide connections that a neuron harbors to its degree within the neuropeptide network (Figure 8D, left panel), with some but not all neuropeptide hubs showing a high diversity of autocrine signaling. In contrast, weak to no correlations were observed between autocrine connection diversity and the synaptic, gap junction, or monoamine connectomes.

**Figure 8.**
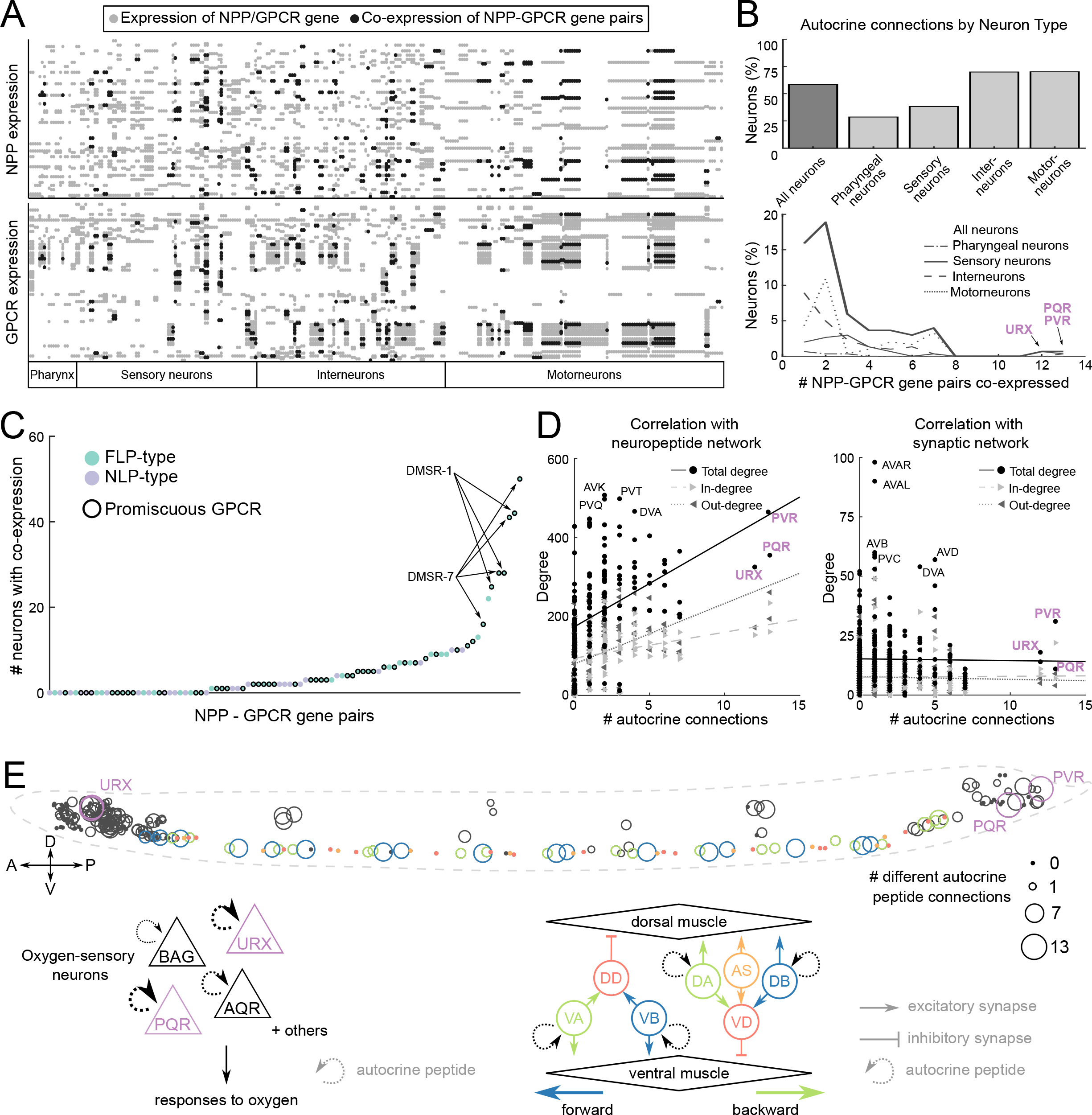
Co-expression between GPCRs and their corresponding neuropeptide ligand(s) facilitates potential autocrine and paracrine signaling. Additional data on putative autocrine signaling is in Supplementary Figure S6. (A) Neuronal expression matrix for both NPP and GPCR genes of the 91 NPP-GPCR pairs. Grey dots represent neuronal expression of only the NPP (upper panel) or GPCR (lower panel) gene, which black dots indicating co-expression of both. (B) Percentage of each neuron type showing peptide autocrine connections (upper panel). The number of different NPP-GPCR pairs being co-expressed in each neuron type is figured in the bottom panel. Neurons showing the highest diversity of different NPP-GPCR pairs being co-expressed are highlighted, including the sensory neurons PQR and URX and the interneuron PVR. (C) Scatter plot showing the number of neurons with co-expression for each of the 91 NPP-GPCRs. Pairs with promiscuous GPCRs, specially DMSR-1 and DMSR-7 are co-expressed in many different neurons, highlighting their central role in autocrine neuropeptidergic signaling. (D) Correlation between number of autocrine connections and neuropeptide (left) or synaptic (right) degree for each of the 302 neurons. Point shapes indicate degree (round), indegree (incoming arrow), outdegree (outgoing arrow). (E) Representation of the autocrine connections in each neuron of the *C. elegans* worm (anterior part of the worm in the left side, posterior in the right, dorsal upper part, and ventral lower part). The size of each cell body indicates the number of autocrine NPP-GPCR pairs expressed in that neuron. Circuits in which autocrine connections are prominent are the locomotion and oxygen sensing circuit. The colors indicate the neurons in the oxygen sensing and locomotion circuits that have the largest number of autocrine connections. Left bottom: oxygen sensing circuits with autocrine connections, URX and PQR are the neurons with the largest number of connections (arrow size indicates number of NPP-GPCR pairs). Right bottom: description of the locomotion system with autocrine connections, VA/VB and DA/DB are excitatory motorneurons that actuate worm crawling and which show many autocrine connections while inhibitory DD/VD and minor excitatory AS motorneurons do not have autocrine connections.

Autocrine connections were especially prevalent in specific circuits in the *C. elegans* nervous system. For example, both URX and PQR, which have among the highest diversity of autocrine signaling, are key sensory neurons that mediate *C. elegans’* responses to aversive O_2_ levels^80–82;^ other O_2_ sensory neurons (AQR and BAG) also co-express ligand-receptor pairs, though to a lesser degree (Figure 8E, bottom left). As O_2_ sensing neurons tonically signal ambient O_2_ concentrations^83^, autocrine signaling may play a stabilizing role in maintaining the homeostasis of neuronal activity. Autocrine neuropeptide signals are also prevalent in the neuromuscular circuit of the ventral nerve cord (VNC – Figure 8F, bottom right) which mediates locomotion^84^. These autocrine loops are primarily mediated by eight RFamide-related neuropeptides that activate two promiscuous receptors, DMSR-1 and DMSR-7, although other receptors such as NPR-5 are also involved (Supplementary Figure 6F). Interestingly, autocrine signaling is restricted to the excitatory A- and B-class VNC motorneurons which drive backward and forward locomotion, respectively^84^; in contrast, both D-type inhibitory motor neurons and the excitatory AS motorneurons, which play modulatory roles in both forward and backward crawling^85^, do not co-express any neuropeptide-GPCR pairs. It should be noted that some autocrine peptidergic pathways are shared between neighboring excitatory motorneurons in the VNC, raising the possibility that they might coordinate activity and/or neurosecretion across excitatory motorneurons in a paracrine manner^86–89^. Heterogeneity in autocrine neuropeptide pathways within the A-and B-class motorneurons (Supplementary Figure 6F) hints at heterogeneity in the contribution of individual motorneurons in this process.

## Discussion

### Neuropeptide signaling forms a complex wireless network

Neuropeptide signaling has long been recognized as critical to brain function, yet despite recent advances in connectomics, the structures of neuropeptidergic signaling networks are largely uncharacterized. We have generated a draft neuropeptide connectome by integrating information from three datasets: a biochemical deorphanization screen matching neuropeptidergic GPCRs with their ligands, a scRNAseq dataset characterizing the expression patterns of all peptide and receptor genes at the single neuron level, and an anatomical dataset defining the morphologies and neuron-neuron contacts of all *C. elegans* neurons. The resulting connectome is remarkably dense; even a short-range network with the most conservative assumptions about the spatial diffusion of neuropeptides was more than ten-fold denser than the previously mapped synaptic connectome of *C. elegans*. As we have also used conservative assumptions regarding gene expression thresholds and ligand-receptor affinities, and since approximately half the putative neuropeptide-activated GPCRs remain to be deorphanized, the actual neuropeptide connectome is almost certainly even more dense than the draft network described here. However, based on sensitivity analysis it seems clear that the overall structure and topological features of the neuropeptide connectivity are likely to hold even when more connections are added to the network (Supplementary Figure 4A-B).

A particularly salient feature of the neuropeptide connectome is its decentralized topology, which contrasts sharply with the more centralized structure of wired neural connectomes. Synaptic connectomes from worms to humans are characterized by a core of high-degree hubs, which are interconnected to form a rich club. This rich club, which in *C. elegans* consists of 11 premotor interneurons, occupies a central position in the connectome, connecting local modules and coordinating their activity. Rich clubs have been previously linked to important functional properties; for example, in *C. elegans* the synaptic rich club has been shown to be involved in global brain states related to locomotion^55, 74^, and in *Drosophila melanogaster* the synaptic rich club constitutes the sensorimotor integrative center of the organism^90^. The gap junction and monoamine connectomes of *C. elegans* likewise contain relatively small rich clubs of less than 20 neurons of relatively high degree. In contrast, the neuropeptide connectome contains a rich club of (in the short-range network) 156 neurons, more than half the neurons in the entire nervous system. The neurons in this neuropeptidergic rich club are extremely well-connected to each other as well as to the rest of the nervous system; all have a degree greater than 203 (in the mid-range network degree of 251), meaning they both send and receive direct connections from at least 40% of all neurons. This remarkable decentralization may be a feature of neuropeptide signaling networks in other organisms^43, 44, 91^.

### Implications of neuropeptide network structure for neuronal computation

The decentralized structure of the neuropeptidergic connectome implies that the strategies it uses for computation and information flow may differ significantly from those employed by the more centrally-organized synaptic network. Intriguingly, both nematode and mammalian neuronal types appear to express nearly unique combinations of neuropeptides and receptors, serving as a molecular bar code for neural identity^40, 43^. Thus, the source of a signal may be encoded by the precise combination of peptides released by the sending neuron. The dynamics of neuropeptide release are also likely to be critical to the information conveyed by peptidergic signals. For example, acute release of FLP-20 peptides by mechanosensory neurons has been shown to trigger short-term sensory locomotor arousal in response to touch stimulation, while chronic release of FLP-20 peptides from the same neurons mediates long-term cross-modal plasticity in olfactory circuits when the sense of touch is lost^14^. Future studies of such mechanisms are likely to provide general insights into how neuropeptide networks encode information in the brain.

Despite its highly dense connectivity, the core of the neuropeptide connectome exhibited a clear substructure. When input patterns were analyzed using dimensionality reduction methods, the neurons in the network core could be divided into three clear groups, one containing mainly pharyngeal and sensory neurons, a second containing the highest degree interneurons, and a third containing many motorneurons. These three groups are themselves connected in a defined pattern, with the second group linking the first and third groups which show few direct links with each other. This organization contrasts in interesting ways with the organization of many synaptic networks, in which peripheral neurons form modules with high internal connectivity that connect to each other through the hubs of the rich club core. In the neuropeptide connectome the core itself exhibits a clear meso-scale organization, forming 3 distinct groups of neurons defined not by unusually high intra-group connectivity, but by intra-group similarity of incoming and outgoing connections patterns. This is in line with recent work showing the existence of different types of meso-scale organization^77^ and is also reminiscent of stochastic block modelling approaches previously applied to the *C. elegans* synaptic connectome^52^. It will be interesting to see if neuromodulatory and synaptic networks from other nervous systems show a similar diversity on meso-scale structure. We expect that looking beyond classical modules is likely to become increasingly important as we begin to look at cellular-scale connectomes of larger organisms, with classes of sensory neurons that are not interconnected but have similar connectivity profiles and perform similar functions.

### Identification of neuropeptide signaling hubs

Although the rich club of the neuropeptide connectome is extensive and encompasses a large portion of the *C. elegans* nervous system, some neurons within this group are of particularly high degree and therefore may play key roles in neuromodulatory signaling. Perhaps not surprisingly, the premotor neurons of the synaptic rich club are also highly connected in the neuropeptide connectome; all 11 of these neurons show a neuropeptide degree above 400 in the short-range network. Since these neurons are known to play important roles in driving global brain states^14, 74, 76^ it is logical that they would also be important targets of neuromodulatory control. In addition, the neuropeptide connectome contains 6 neurons whose neuropeptide degree is higher than any of the synaptic rich club (degree > 490). Three of these neurons (PVT and the pair of AVK neurons) are specialized peptidergic neurons that express no classical neurotransmitter or monoamine and have been linked to arousal and sleep-like behaviors^14, 92, 93^. Specialized neuropeptidergic neurons have also been described in other organisms, and in mice they have been linked to global behaviors such as fear^94^.

The other peptidergic hubs (PVR and the pair of PVQ neurons) are tail neurons that extend long processes to the nerve ring, and in the case of the PVQs (as well as PVT) these processes are rich in dense-core vesicles^48, 65, 66^. Since AVK also has a long process (extended from its cell body in the head through the nerve ring and the ventral cord to the tail) these neurons may be morphologically specialized for local release of peptides throughout the nervous system. While AVK is known to play roles in arousal and motor control, the functions of the remaining neuropeptide hubs (or of other neurons of high neuropeptidergic degree such as PVN and PVP) are largely uncharacterized. Given their importance in the neuropeptide signaling network, it will be interesting in the future to explore the roles of these neurons in the control of behavioral states.

It is interesting to note that even in the long-range network, where no spatial restrictions on neuropeptide diffusion are imposed, the six short and mid-range hub neurons retain their central importance in the network. This confirms that the high degree of the hubs in the short- and mid-range networks is not merely an artifact of the spatial conditions we imposed but rather is a consequence of expressing key combinations of neuropeptides and receptors. Notably, the long-range network contains additional high-degree nodes that are not hubs in the short-and mid-range networks, in particular the oxygen-sensing neurons URX, AQR, PQR and BDU and the tail motorneurons DA09 and VA12. One might speculate that these neurons could in fact engage in long-range neuroendocrine signaling, and that the long-process morphology of the short-and mid-range hubs might allow them to carry out neuropeptide modulation on a finer temporal or spatial scale. In the future, these questions may be addressable using *in vivo* probes for neuropeptide receptor signaling^95, 96^.

### Neuropeptide signaling links specific components of the nervous system

In addition to these high degree nodes, the neuropeptide connectome also contains edges of unusually high weight, representing neuron pairs linked by multiple neuropeptide-signaling pathways. While many neuron pairs in the network are connected by a single neuropeptide pathway, 17 pairs of neurons are linked by 15 or more different neuropeptide-receptor couples. Most of these extremely high-weight edges involve connections between the oxygen-sensing neurons PQR or URXL/R and the motor circuit (either the DB motorneurons or the AVD premotor interneurons). Why might oxygen-sensing neurons participate in so many complex neuromodulatory interactions? These neurons strongly influence locomotor states and their tonic responses to ambient oxygen are influenced by experience and other sensory cues. Complex neuropeptide signaling may allow feedback between the oxygen sensors and motorneurons to fine-tune the activity of this circuit across time and space.

The oxygen-sensing neurons and the motor circuit were also found to be important sites for autocrine signaling, in which a neuropeptide receptor and its ligand are co-expressed in the same neuron. Autocrine peptide signaling is ubiquitous in other brains as well^97, 98^, suggesting it supports important aspects of neural function. For example, autocrine neuropeptide pathways are known to mediate cell-autonomous feedback^99–104^, and are therefore often suggested to maintain neuronal homeostasis; this may be particularly important to regulate the activity of tonically signaling neurons, such as the oxygen-sensing neurons. In the motor circuit, ostensibly autocrine signaling may play a further role by coordinating the activities of neighboring neurons with similar gene expression patterns. Although proprioceptive and electrical coupling between adjacent motorneurons generates waves of muscle contraction over adjacent body regions^88^, autocrine pathways also have the capacity to coordinate physiology across neighboring cells^105, 106^. Indeed, mutants for several of the neuropeptide ligands that act in motorneuron autocrine pathways have been shown to have locomotion defects^107, 108^, suggesting an important role for these connections in patterning locomotor behavior.

We also observe a critical role for neuropeptide signaling in linking disconnected components of the nervous system to the broader network. In particular, the pharyngeal neurons, which form a wired network analogous (if not homologous) to the vertebrate’s enteric nervous system, and the CAN neurons, whose processes lie in the excretory canal-associated nerve, are virtually completely unconnected to the somatic synaptic and gap junction connectomes yet are extensively integrated into the neuropeptide connectome. Both CAN and the pharyngeal neurons express multiple GPCRs whose ligands are expressed exclusively outside the pharynx or canal-associated nerve, indicating that these neurons are regulated by peptides released from physically-unconnected processes. Despite their disconnection from the wired connectomes, the pharynx and CAN neurons carry out essential physiological functions; indeed, CAN and the pharyngeal neuron M4 are the only neurons whose ablation is lethal to the animal^109, 110^. This provides a means for communication between the wider nervous system and these isolated but biologically critical neurons.

### Prospects for future mapping of wireless brain connectomes

We have described here the draft neuropeptide connectome of *C. elegans*, an animal with 302 neurons. In the future, we plan to refine this connectome, for example by expanding its scope through deorphanization of neuropeptide receptors whose ligands are currently unknown. Differential posttranscriptional processing may also lead to the synthesis of different peptides and receptors in neurons that express the same gene, and alternative patterns of gene expression during development or in response to environmental cues may likewise alter the structure and function of neuropeptide signaling networks. Moreover, non-neuronal cells function as both senders and receivers of neuropeptide signals; with the use of reporter lines for peptides and receptors it should be possible in the future to incorporate these cells into the neuropeptide connectome and observe plasticity in the network. We likewise plan to integrate functional information into the connectome map; for example, in vitro experiments could identify the G-protein pathways downstream of individual receptors, and *in vivo* sensors could provide empirical data on the spatial scope of neuropeptide signaling pathways.

Together these data will facilitate functional modeling of neuropeptidergic circuits revealed in the connectome maps.

In principle, the approaches described here should also be applicable to mapping the neuropeptide signaling networks of animals with much larger brains. Single-cell RNAseq data from different vertebrate animals indicates that, as in *C. elegans*, most neurons express at least one neuropeptide precursor and at least one neuropeptide receptor, facilitating dense and potentially decentralized networks^18, 39, 43, 44^. Although it is currently not possible to precisely link gene expression clusters with individual neurons in brain circuits, the use of targeted reporters should eventually make it possible to relate neuropeptide and receptor expression to increasingly detailed synaptic connectome maps in flies and mice. Basic mechanisms of neuropeptide signaling are shared in all animals, from nematodes to mammals: neuropeptides are released from dense core vesicles and diffuse across space to neurons unconnected to the releasing cell by wired synapses. And although the *C. elegans* nervous system is anatomically small, at the molecular level its neuropeptide systems are highly complex and show significant homology to other animals. Thus, the neuropeptide connectome of *C. elegans* may serve as a prototype to unravel general principles of neuromodulatory network structure that also apply to much larger brains.^34, 38, 35,36,37,18^

## Methods

### Reporter Transgenic Strains

*CRISPR based constructs*: Thirty-two transcriptional C-terminal GFP reporters of neuropeptides and neuropeptide receptors were created where GFP was recombineered in the last coding exon of the gene of interest. Of the 17 total neuropeptide precursor genes that we made reporters for, 8 had only a T2A::3xNLS::GFP tag, 1 had a T2A::GFP::H2B tag, 6 had both T2A::3xNLS::GFP and SL2::GFP::H2B tags, and 2 had only an SL2::GFP::H2B tag (Supplementary Table 1). For the GPCR receptors, 8 gene expression reporters were made all with SL2::GFP::H2B tags (Supplementary Table 2). Most of these constructs were made by SUNY Biotech.

### Reporter Analysis

Young adult animals were mounted on 5% agarose pads and immobilized with 100 mM sodium azide and imaged on a Zeiss LSM880 using a 40X objective lens. GFP expression reporters were identified at single neuron resolution as described^111^. GFP reporter expression of these constructs, as reported in Table S1-2 were noted using three categories: moderate to high expression, low and variable expression, and no detected expression. Additionally, we compared our GFP reporter expression data to single-cell RNA-seq expression (scRNAseq) data from the CeNGEN project using their standard thresholds (4 being the most stringent, 1 being the least stringent, and blank be unfiltered)^40^.

In our analysis, for each gene and each CeNGEN threshold, we tallied 1) the number of neurons that showed GFP expression but not scRNA expression, 2) the number of neurons showing both GFP expression and scRNA expression, and 3) the number of neurons that showed scRNA expression but no GFP expression (Table S3).

Based on the results of this analysis threshold 4, although in some occasions too conservative, had the best correlation between GFP reporter and CeNGEN scRNAseq data expression per neuron for the tested NPP and GPCR genes.

### Synaptic and gap junction networks

The synaptic and gap junction networks used in this work were based on the full hermaphrodite *C*. *elegans* connectome, containing all 302 neurons. This network was composed from the somatic connectome^48^, updated and released by the Chklovskii lab^47^; and the pharyngeal network of Albertson and Thomson^65^, made available by the Cybernetic *Caenorhabditis elegans* Program (CCeP). The functional classifications referred to in the text (i.e., *sensory neuron*, *interneuron*, *motorneuron*) are based on the classification scheme used in WormAtlas^112^. When there is double or triple classification in nerve ring neurons that can be sensory neurons, interneurons, or motorneurons, the Zhen lab classification^66^ was used to select one neuron type. URB left and right neurons are the only ones in which the WormAtlas and the Zhen lab classification diverge, leading us to classify them as sensory neuron following the later most recent classification. DB neurons are identified as motorneurons although WormAtlas indicates that these could also be interneurons^112^. The gap junction network was modelled as an undirected network with bidirectional electrical synapses; note however that some gap junctions might be rectifying and thus exhibit directionality. In the synaptic network reciprocal connections between nodes are considered as two separate unidirectional connections.

### Monoamine network construction

The monoamine network used in this work was made following the same procedure as in (Bentley et al., 2016). The monoamine expression for the 302 neurons comes from the neurotransmitter atlas of *C. elegans*^75^ and receptor expression for the 302 neurons comes from the single-cell expression data from the CeNGEN project (https://www.cengen.org)^40^. We used the expression data at CeNGEN threshold 4. The interactions between ligand and receptors were previously described^45^. The adjacency matrix was built using a binary version of the expression data for the 302 neurons. For a given point A^M^(i,j) and for a given monoamine receptor pair M the connection between two neurons is defined by A^M^(i,j) = Mon^M^(i,j) x Receptor^M^(i,j). Each monoamine receptor interaction forms an individual binary network. To get the overall monoamines network we add each individual monoamine receptor network resulting in a weighted network where the weight indicates the number of monoamine receptor pairs that connect two nodes. Reciprocal connections between nodes are considered as two separate unidirectional connections.

### Neuropeptide network construction

The neuropeptide network used in this work was made using a similar approach to that used for the monoamines. The interactions between ligands and receptors were identified using a large-scale *in vitro* reverse pharmacology pipeline in which over 87% of the predicted peptide GPCRs were challenged with FMRFamide related peptides (FLP) and non-insulin non-FLP like peptides (NLP)^64^. Neuropeptide precursor and GPCR gene expression for the 302 neurons was extracted from the single-cell transcriptome data of the CeNGEN project (https://www.cengen.org^40^). We used the expression data at CeNGEN threshold 4. The adjacency matrix was built using a binary version of the expression data for the 302 neurons. For a given point A^N^(i,j) and for a given neuropeptide receptor pair N the connection between two neurons is defined by A^N^(i,j) = NPP^N^(i,j) x GPCR^N^(i,j). Each neuropeptide receptor interaction forms an individual binary network. To get the overall neuropeptides network we add each individual neuropeptide receptor network resulting in a weighted network where the weight indicates the number of neuropeptide receptor pairs that connect two nodes. Reciprocal connections between nodes are considered as two separate unidirectional connections.

### Neuropeptide network spatial constraining

Neuropeptidergic networks were locally thresholded to filter out connections between neurons that were anatomically far from each other. The anatomical EM data was obtained from The Mind of the worm (https://www.wormatlas.org/MoW_built0.92/MoW.html) and other literature^48, 65, 66^. This data was used to create a table of locations for each neuronal process, identifying 27 different neuronal process bundles in the *C. elegans* nervous system as previously defined^48^. This classification was then used to filter out neuropeptidergic connections based on putative signaling ranges. The stringent short-range thresholding allows connections only in between neuronal processes that are in the same process bundle and the pharynx is a separated system were all connections are allowed in between pharyngeal neurons only. The mid-range stringency thresholding allows connections between neurons with neuronal processes in the same anatomical area: head (including pharynx and the ventral cord neurons that are in the ventral ganglion), midbody and tail. And in the unthresholded system all neuropeptidergic connections are allowed.

### Topological network measures

Edge counts, adjacency matrices and reducibility clusters were all computed using binary directed versions of the networks. The same networks, excluding self-connections (i.e. setting all diagonal elements to 0), were used to compute all other measures.

Network measures are compared to 100 null model networks generated using the degree-preserving edge swap procedure from the Brain Connectivity Toolbox for MATLAB^113^. This is performed by selecting a pair of edges (*A*→*B*) (*C*→*D*) and swapping them to give (*A*→*D*)(*C*→*B*). If the resulting edges already exist in the network, another pair of edges is selected instead. Each edge was swapped 10 times to ensure full randomization.

### Degree

Degree is the number of edges connected to a given node. Indegree is the number of incoming connections connected to a given node and outdegree is the number of outgoing connections.

### Density

Density *d* is the fraction of present connections *K* to possible connections between the given nodes *N*: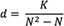

### Clustering coefficient

Transitivity defines the ratio of triangles to triplets in the network (where a triple is a single node with edges running to an unordered pair of others, and a triangle is a fully-connected triple). For a directed network, this is equivalent to: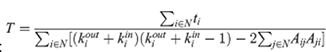 where *A* is the adjacency matrix, *N* is the number of nodes, *k^out^* and *k^in^* are the out-degree and in-degree, and *t_i_* is the number of triangles around a node: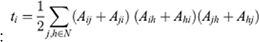

### Reciprocity

Reciprocity is the fraction of reciprocal edges in the network: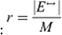where *M* is the number of edges, and |*E*^↔^| is the number of reciprocal edges: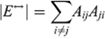

### Rich-club coefficient

The rich-club coefficient measures the tendency for high-degree nodes in a network to form highly interconnected communities^55^. These communities can be identified by creating subnetworks for each degree level *k* and removing nodes with a degree ≤ *k*. Then the rich-club coefficient Φ(*k*) for each subnetwork is defined as the ratio of connections in the subnetwork *M_k_* to the number of maximum possible connections. For a directed network with no self-connections, where *N_k_* is the number of remaining nodes, this is given by: 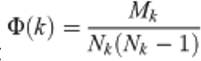

Thus, a fully connected subnetwork at a given degree *k* has a rich-club coefficient Φ(*k*) = 1. We normalize the rich-club coefficient by calculating: 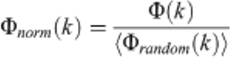were ˂Φ_random_(*k)* ˃ is the average value of the rich-club coefficient across random networks.

A rich-club exists when Φ*_norm_*(*k*) ≥ 1, but in order to get a clear threshold range we use a probabilistic approach. The threshold range of the rich-club is defined by Φ*_norm_*(*k*) ≥ 1 + 1*σ*, where *σ* is the Standard Deviation of Φ*_random_*(*k*) for the 100 random networks.

### Dimensionality reduction analysis

t-SNE is an algorithm for dimensionality reduction that facilitates visualizing high dimensionality data. The analysis described here was performed using the MATLAB *t-sne* function on the adjacency network of connections. The neuropeptide dimension was reduced, and clustering was performed based on the pattern of connections due to receptor expression. Different distant measures were tested to confirm the clustering: Euclidean distance, Chebychev distance, cosine distance and Mahalanobis distance.

### Co-expression analysis

The signaling networks used in this work represent connections between co-occurring genes. Nodes are defined as pairs of neuropeptide precursor and GPCR genes that co-occur more than expected by chance as measured by a Fisher’s exact test^114^ with FDR (false positive rate) correction^115^. The 2x2 contingency table for the Fisher’s exact test contains the number of neurons for which both genes co-occur and the number of neurons in which each NPP and GPCR gene is expressed without co-occurring with the second gene. Thus, the Fisher’s test is defined as:

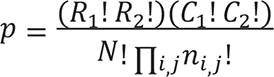

Where R_1_ and R_2_ are the row sums, C_1_ and C_2_ are the column sums, *N* is the total number of observations in the contingency table, and n_ij_ is the value in the *i*th row and *j*th column of the table

Interactions between nodes are defined by the receptor-ligand interactions that the co-occurring genes have with genes that co-occur in another node. The interactions between ligand and receptors were identified using a large-scale *in vitro* reverse pharmacology pipeline^64^.

### Software

Network measures were computed in MATLAB (v9.8.0.1323502 (R2020a), The MathWorks Inc. Natick, MA) using the Brain Connectivity Toolbox^113^ (v2019-03-03) and the MATLAB/Octave Networks Toolbox^116^. Clustering and visualization of multilayer plots was performed using MuxViz^117^. And additional network visualizations were created using Cytoscape^118^.

### Lead contact

Requests for resources should be directed to the Lead Contact William Schafer

### Data and software availability

The scRNAseq data are available at www.cengen.org. The biochemical deorphanization data are available at https://worm.peptide-gpcr.org/project/neuropeptides/flps/ ^64^. Analysis code is available at https://github.com/LidiaRipollSanchez/Neuropeptide-Connectome.git

### Nematode Strains

The strains used in this study are listed in Supplementary Tables 1 and 2.

## Acknowledgements

This work was funded by grants from NIH (R01NS110391, to OH, WRS and IB), the MRC (MC-A023-5PB91, to WRS), the Wellcome Trust (WT103784MA, to WRS), MQ: Transforming Mental Health (MQF17_24, to PV) the ERC (950328, to IB), HHMI (to OH), and the Simons Foundation (855199, to RF). This research was also supported in part by the NIHR Cambridge Biomedical Research Centre (BRC-1215-20014) and NIHR Applied Research Centre. The views expressed are those of the author(s) and not necessarily those of any funding bodies. For the purpose of open access, the authors have applied a Creative Commons Attribution (CC BY) license to any Author Accepted Manuscript version arising from this submission.

We’d like to thank the LMB Visual Aids and Scientific Computing teams for their support, especially Johanna Westmorland for figure 3 design. We thank Tom Shimizu, Erik Larsson Lekholm, Mark Moyle, Caroline Nettekoven, Sarah Morgan, Albert Cardona, Ed Bullmore, Amy Courtney and Mei Zhen for productive discussions. We thank WormAtlas, OpenWorm, Nemanode and WormBase for their resources.

## Author contributions

L.R.S., W.R.S., P.E.V., J.W. and I.B. designed research. L.R.S. and J.W. performed analysis. H.S. and R.F. made strains and interpreted images. H.S., R.F., S.R.T., A.W., M.H., D.M.M., O.H. contributed with data and discussion. L.R.S., W.R.S., P.E.V., J.W. and I.B. contributed to writing the paper. All authors reviewed and approved the manuscript.

---

**Supplementary Figure S1.**
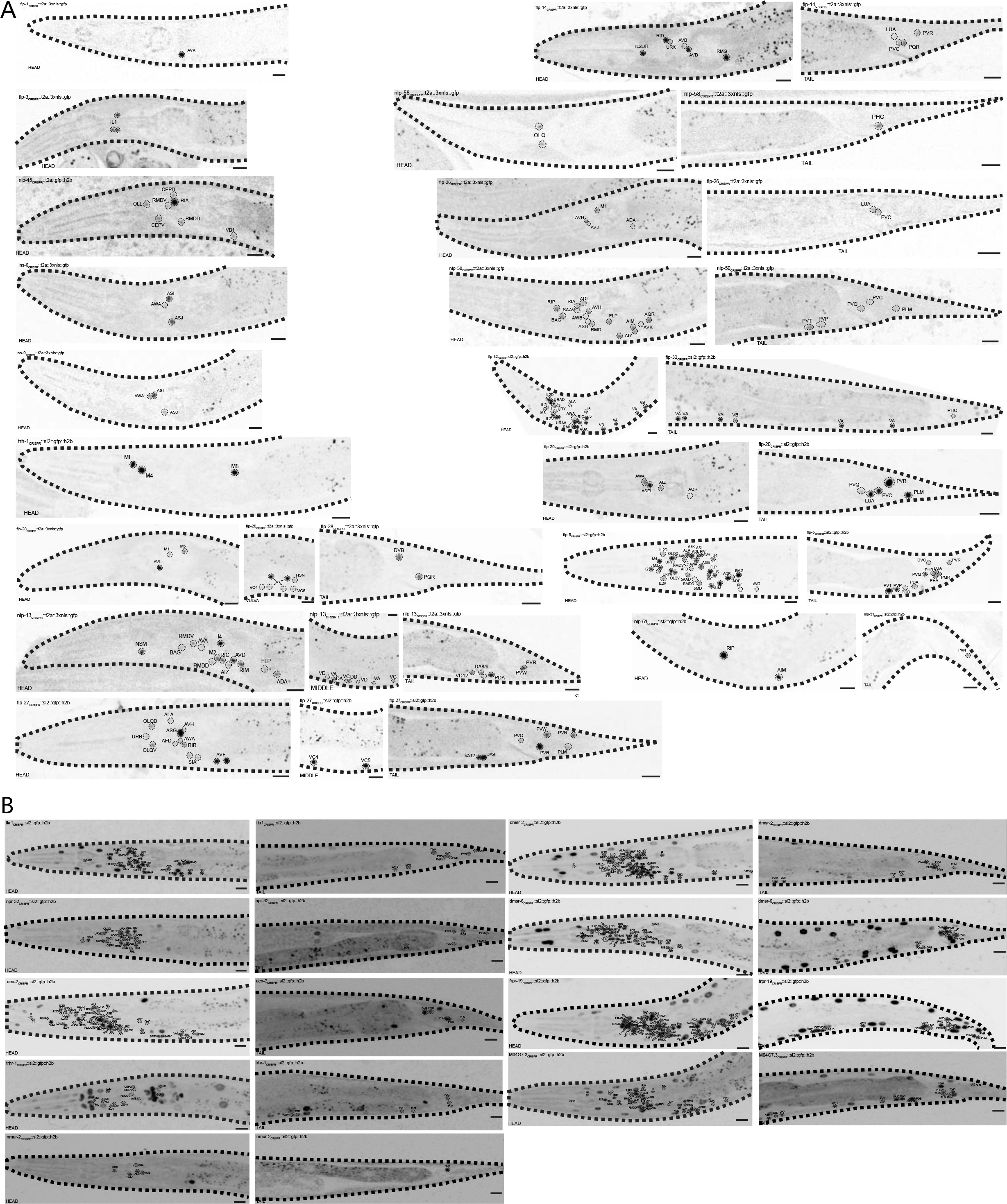
Expression patterns of neuropeptide and GPCR reporters. Cells were identified as described in Figure 2. A) Fluorescent GFP reporters for the expression of 17 representative neuropeptide precursor genes. B) Fluorescent GFP reporters for the expression of 8 representative GPCR precursor genes. Images of the *trhr-1* expression strain show some bleed-through signal that does not represent neuronal expression.

**Supplementary Figure S2.**
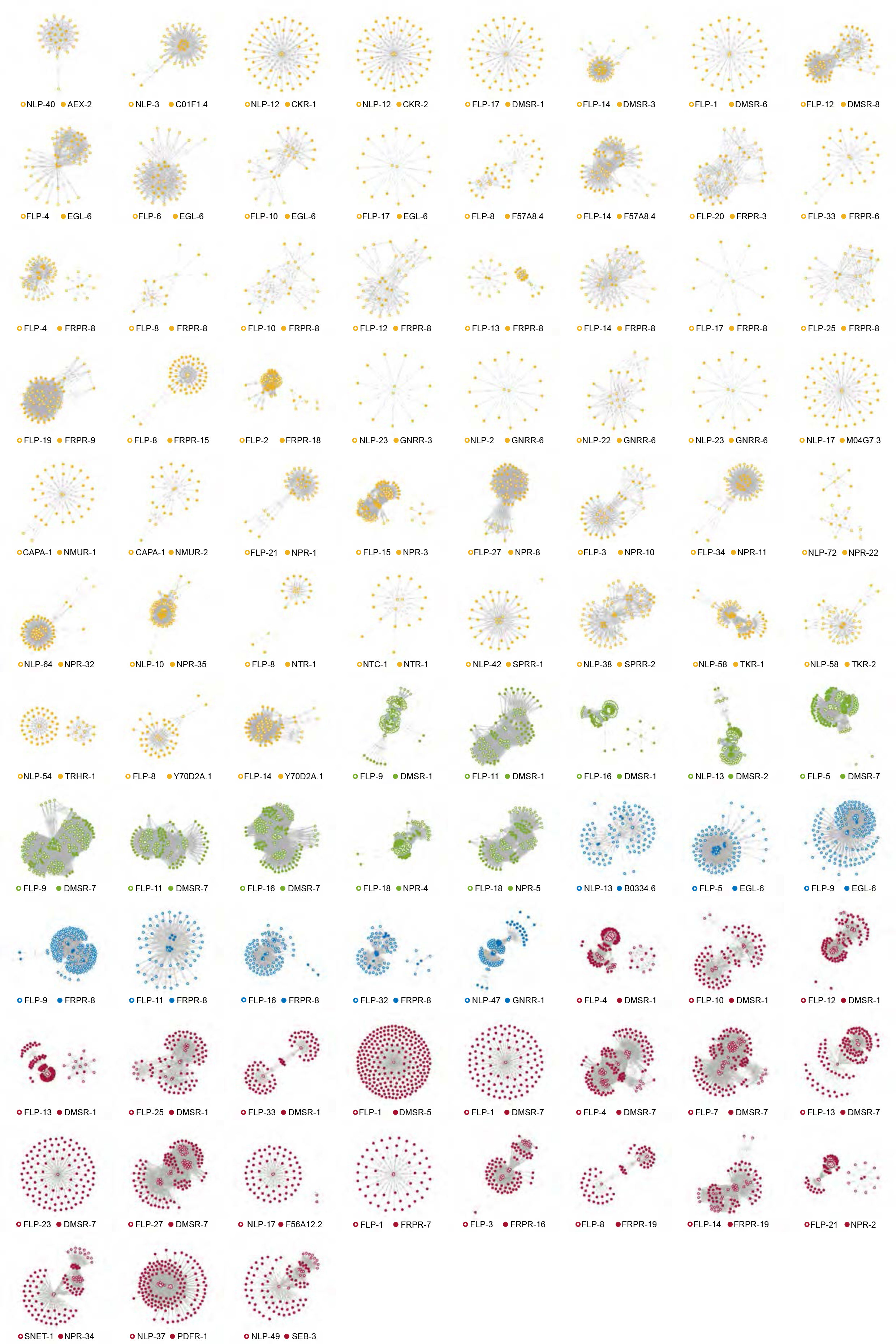
Graphical representations of the 91 NPP-GPCR networks with short-range connections. Colors given by assortativity like Figure 4; yellow indicates local, green pervasive, blue integrative and red promiscuous networks.

**Supplementary Figure S3.**
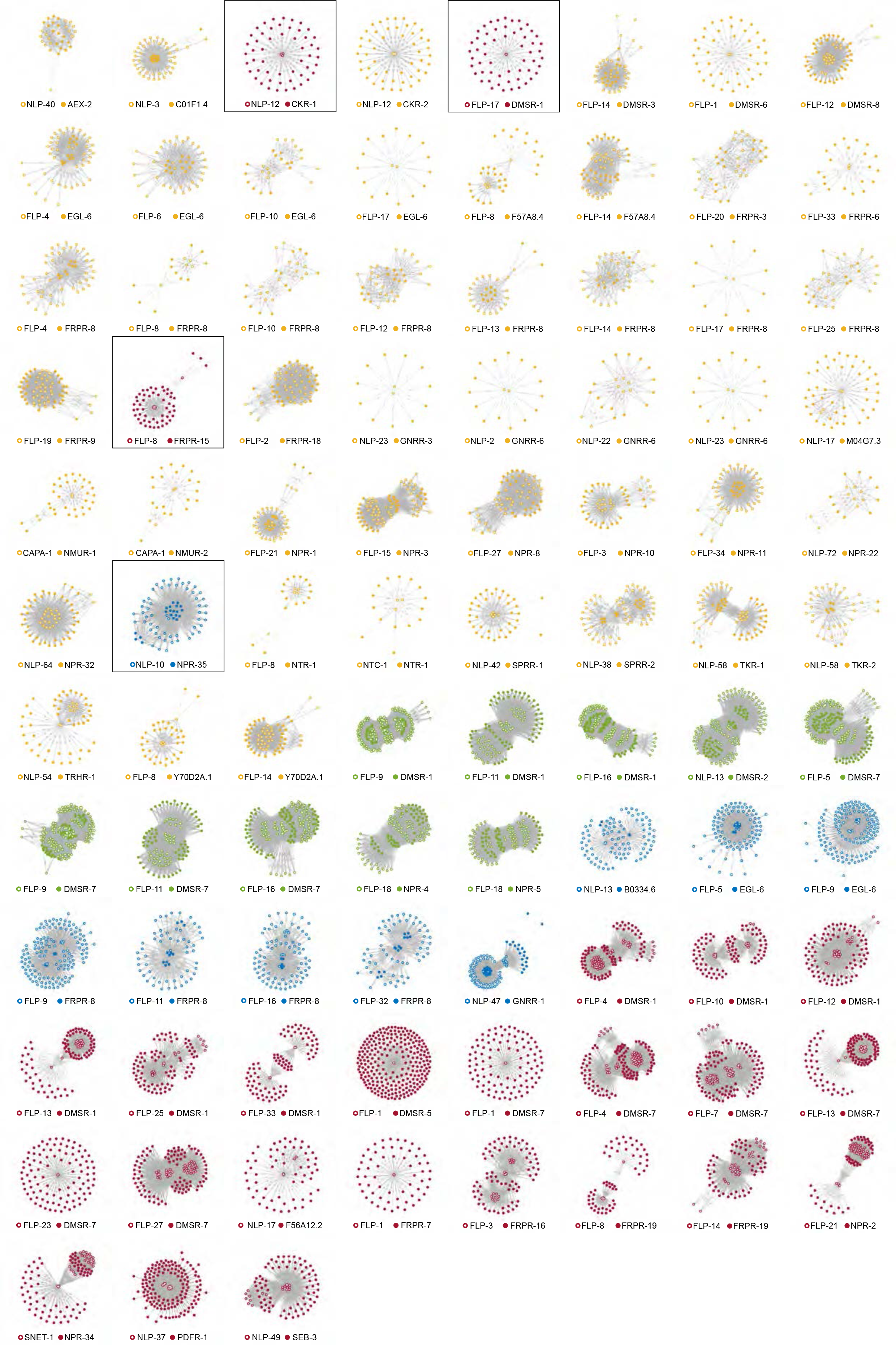
Graphical representations of the 91 NPP-GPCR networks with mid-range connections. Networks are sorted in the same order as in Supplementary Figure 2 for comparison. Colors given by assortativity as defined in Figure 4, yellow indicates local, green pervasive, blue integrative and red promiscuous networks. Networks that change assortativity between short and mid-range spatial models of neuropeptide transmission are highlighted.

**Supplementary Figure 4.**
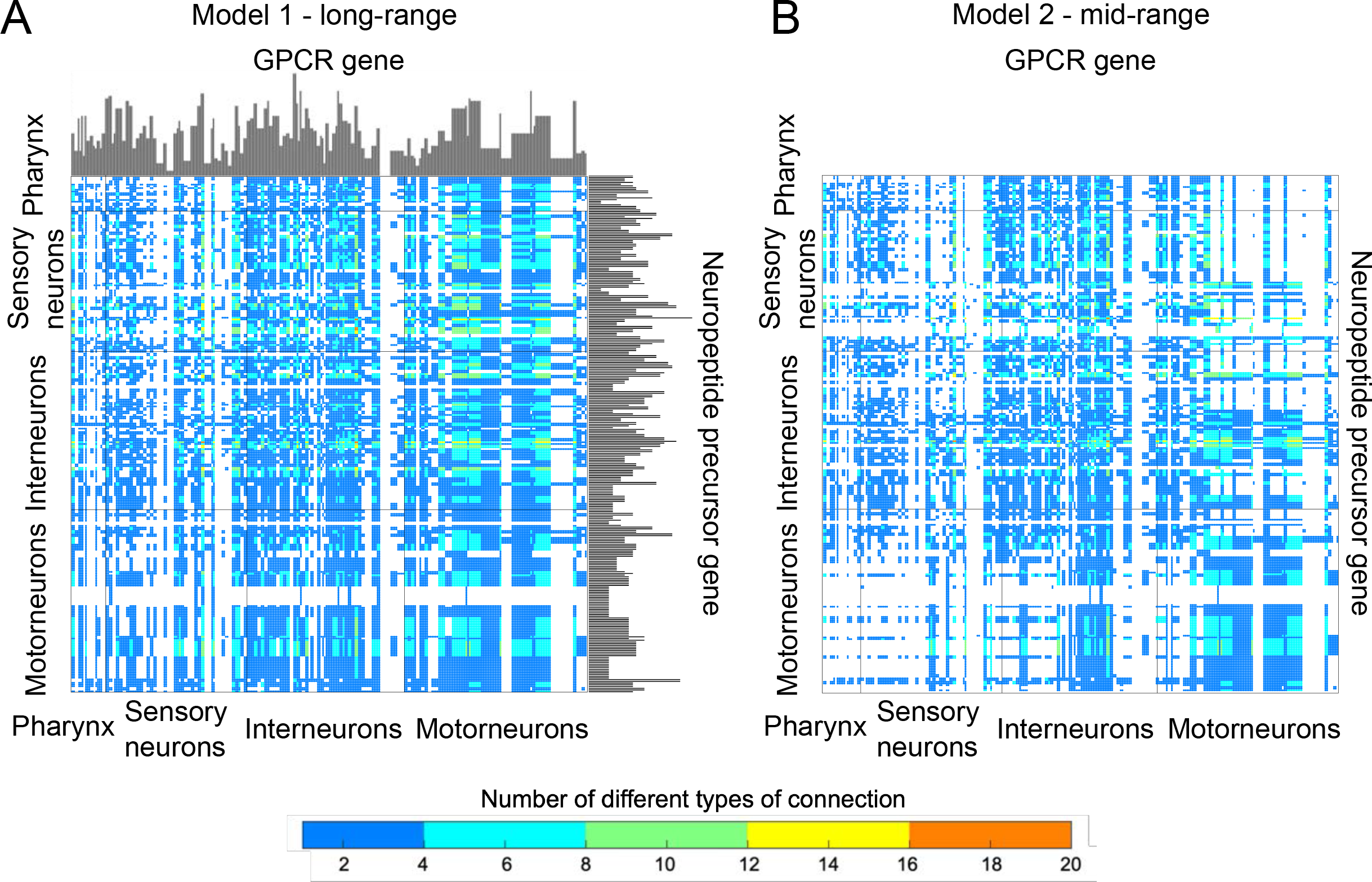
Adjacency matrix representation of the long and mid-range neuropeptide networks. A) Model 1, aggregate matrix of the long-range neuropeptide connections, the histogram lining the matrix indicate the number of GPCR or NPP that the neuron expresses. B) Model 2, aggregate matrix of the mid-range neuropeptide connections.

**Supplementary Figure 5.**
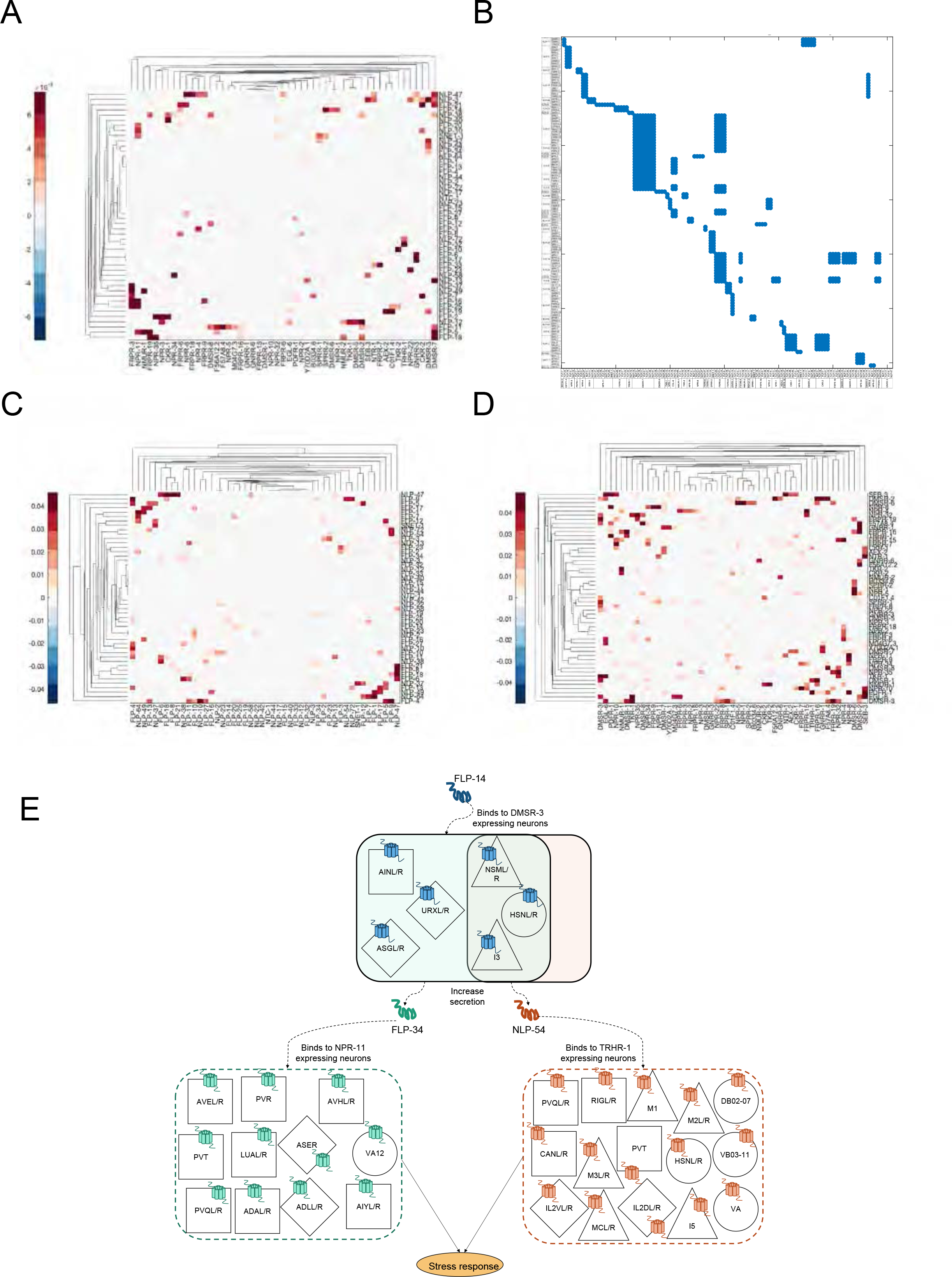
Co-expression analysis of NPP and GPCR genes. A) Hierarchical clustering matrix showing NPP – GPCR pairs with significant co-expression. The y axis indicates GPCR and the x axis indicates NPP expression. The significance of their co-expression is indicated by the legend. B) Adjacency matrix representation of the co-expression network. The nodes represent NPP – GPCR pairs with significant neuronal coexpression, and edges are formed when the NPP of one node is the ligand of the GPCR of another node. C) Hierarchical clustering matrix of the NPP vs NPP co-expression. D) Hierarchical clustering matrix of the GPCR vs. GPCR co-expression. E) Visual representation of one of the possible signaling cascades based on the co-expression network. DMSR-3 co-expresses with FLP-34 and NLP-54 releasing neurons, which means that secretion could be activated by FLP-14 binding to DMSR-3.

**Supplementary Figure 6.**
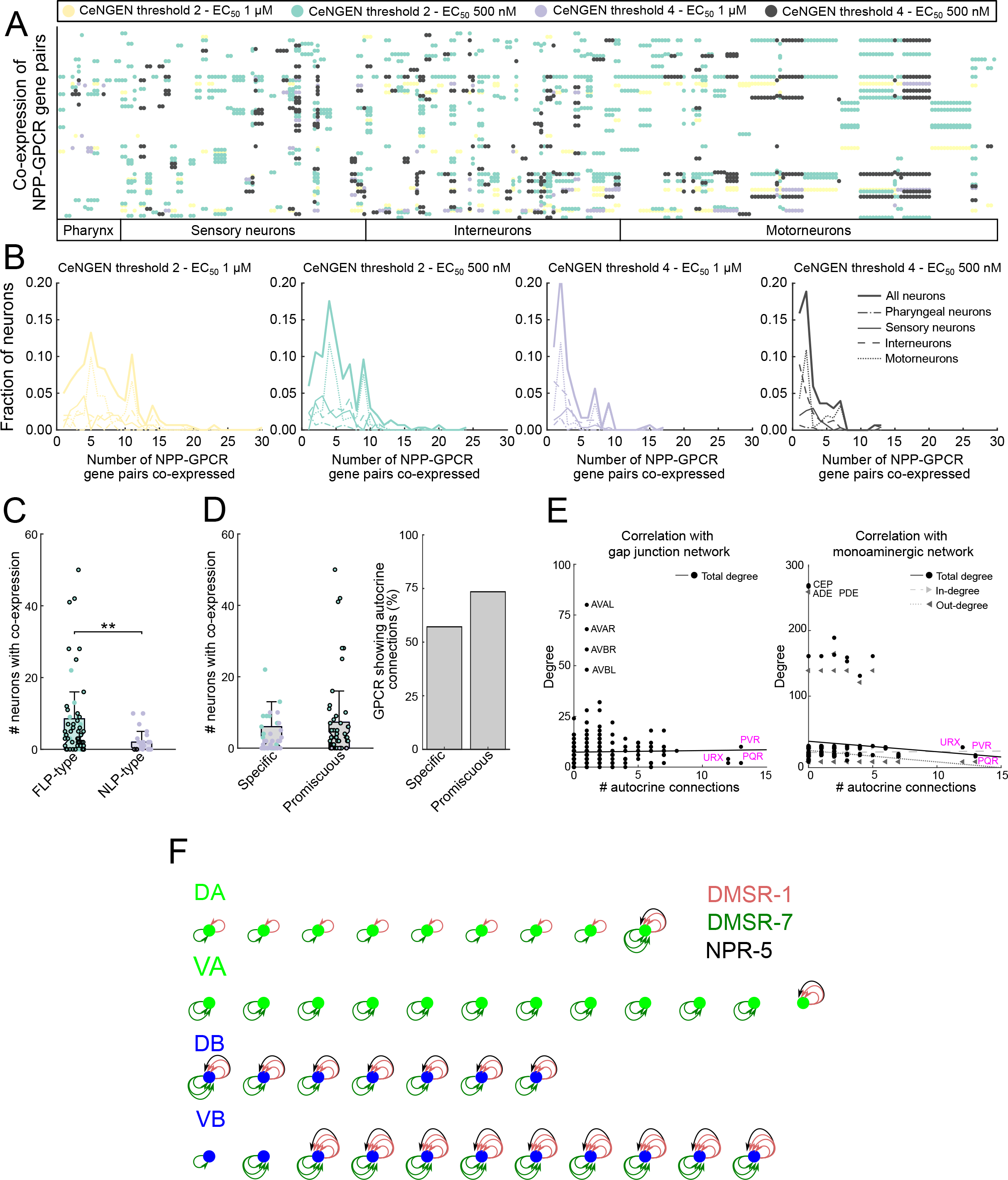
Autocrine connections sensitivity analysis. A) comparison of co-expression of cognate NPP-GPCR pairs in the 4 different combinations of CeNGEN and deorphanization datasets with different stringencies (CeNGEN threshold 2 or 4; EC_50_ threshold 1 µM or 500 nM), each depicted in a different color. B) Number of NPP-GPCR co-expressed per fraction of neurons in these 4 different combinations of datasets. C) Number of neurons with co-expression for the FLP or NLP neuropeptides show that FLP are more prevalent. D) Comparison of number of autocrine connections between specific and promiscuous NPP-GPCR pairs, showing that promiscuous pairs tend to have more autocrine connections. E) Correlation of autocrine connections with gap junction and monoamine networks. F) Number of autocrine connections in each motorneuron for each of the main promiscuous NPP-GPCR pairs present in those neurons.

